# Neuropathological assessment of the olfactory bulb and tract in individuals with COVID-19

**DOI:** 10.1101/2023.12.18.572180

**Authors:** Nathalie A. Lengacher, Julianna J. Tomlinson, Ann-Kristin Jochum, Jonas Franz, Omar Hasan Ali, Lukas Flatz, Wolfram Jochum, Josef Penninger, aSCENT-PD Investigators, Christine Stadelmann-Nessler, John M. Woulfe, Michael G. Schlossmacher

## Abstract

The majority of patients with Parkinson disease (PD) experience a loss in their sense of smell and accumulate insoluble α-synuclein aggregates in their olfactory bulbs (OB). Subjects affected by a SARS-CoV-2-linked illness (COVID-19) frequently experience hyposmia. We previously hypothesized that α-synuclein and tau misprocessing could occur following host responses to microbial triggers. Using semiquantitative measurements of immunohistochemical signals, we examined OB and olfactory tract specimens collected serially at autopsies between 2020 and 2023. Deceased subjects comprised 50 adults, which included COVID19+ patients (n=22), individuals with Lewy body disease (e.g., PD and dementia with Lewy bodies (DLB; n=6)), Alzheimer disease (AD; n=3), other non-synucleinopathy-linked degenerative diseases (e.g., progressive supranuclear palsy (PSP; n=2) and multisystem atrophy (MSA; n=1)). Further, we included neurologically healthy controls (HCO; n=9) and those with an inflammation-rich brain disorder as neurological controls (NCO; n=7). When probing for inflammatory changes focusing on anterior olfactory nuclei (AON) using anti-CD68 immunostaining, scores were consistently elevated in NCO and AD cases. In contrast, inflammation on average was not significantly altered in COVID19+ patients relative to controls, although anti-CD68 reactivity in their OB and tracts declined with progression in age. Mild-to-moderate increases in phospho-αSyn and phospho-tau signals were detected in the AON of tauopathy-and synucleinopathy-afflicted brains, respectively, consistent with mixed pathology, as described by others. Lastly, when both sides were available for comparison in our case series, we saw no asymmetry in the degree of pathology of the left versus right OB and tracts. We concluded from our autopsy series that after a fatal course of COVID-19, microscopic changes -when present-in the rostral, intracranial portion of the olfactory circuitry generally reflected neurodegenerative processes seen elsewhere in the brain. In general, inflammation correlated best with the degree of Alzheimer’s-linked tauopathy and declined with progression of age in COVID19+ patients.

## Introduction

Parkinson disease (PD) is traditionally characterized clinically by extrapyramidal motor dysfunction and pathologically by the progressive degeneration of dopamine producing neurons in the substantia nigra. Over the past two decades, this nigral-and moto-centric view of PD has undergone a revision with the recognition that the disorder is also characterized by a wide variety of non-motor symptoms referable not only to brain pathology but to the peripheral nervous system as well [1]. Indeed, these symptoms may predate the onset of extrapyramidal motor dysfunction by decades [1]. Accordingly, a histopathological hallmark of typical, late-onset PD, namely intracellular α-synuclein (αSyn) aggregation in the form of Lewy bodies (LBs) and Lewy neurites (LNs), has been described in extra-nigral sites within the brain as well as within peripheral organs, often prior to their emergence in the substantia nigra. Braak and Del Tredici [2] provided pathological evidence for a dynamic spatiotemporal sequence of αSyn pathology in the brain. According to their hypothesis, the OBs and dorsal motor nuclei of the vagus nerve represent the earliest CNS sites in which αSyn aggregation occurs. Implicit in the Del Tredici-Braak model is the phenomenon of ‘spreading’ pathology, whereby -once initiated at those sites-the aggregation process is transmitted trans-synaptically to other nuclei including the substantia nigra, resulting in the well-recognized motor features of the disease [3]. Consistent with this hypothesis, hyposmia is one of the earliest and most frequent non-motor signs of PD [1,4], which accords with the presence of Lewy pathology in the OBs as well as in higher order olfactory structures [2]. As a potential trigger of αSyn aggregation in the olfactory system and enteric nervous system, Braak and Del Tredici invoked an infectious, possibly viral, pathogen [3]. In this context, their hypothesis is compatible with the growing body of evidence implicating microbial encounters as risk factors for neurodegenerative disorders in later years (reviewed in [5]).

Subsequent to its discovery in Wuhan, China in late 2019, coronavirus-linked disease 2019 (COVID-19), caused by severe acute respiratory syndrome coronavirus-2 (SARS-CoV-2), spread rapidly throughout the world, culminating in a global pandemic that imposed substantial human suffering and loss of life as well as unprecedented social and economic consequences [6]. Olfactory dysfunction in the form of anosmia, hyposmia, and parosmia were described as a common symptom during the pandemic, affecting 30–70% of all patients with confirmed COVID-19 [7]. Abnormal olfaction usually occurs early in the course of infection, presenting as the first symptom in approximately 12% of all patients [8]. Although most patients recovered their sense of smell gradually, often within 3 to 4 weeks [9,10], some suffered from persistent impairment, suggesting severe and/or permanent damage to components within the olfactory circuitry [9]. Several possible processes have been invoked as mechanisms underlying olfactory dysfunction in COVID-19, including direct infection of olfactory sensory neurons (OSNs) in the nasal cavity [11] as well as primary infection of sustentacular cells with secondary injury to OSNs. In accordance with the latter, several investigators have noted the absence of the SARS-CoV-2 target receptor, angiotensin converting enzyme-2, on neuronal cells, and instead proposed infection and damage to non-neuronal support cells in the olfactory epithelium [12,13]. Regardless of the cellular target, there is histopathological evidence that infection by SARS-CoV-2 can confer effects onto the CNS in the form of local inflammation, axonal pathology and microvascular changes in the OB and olfactory tracts (OT) [14].

Moreover, a recent population-based neuroimaging study in survivors of a SARS-CoV-2 infection revealed chronically reduced volumes of olfaction circuitry-associated areas in the brain. It provided evidence that the potential impact of SARS-CoV-2 on scent processing extends beyond the OT to impose lasting neurodegenerative and/or remodeling changes in the central olfactory circuitry [15].

That COVID-19 and PD share olfactory dysfunction as an early clinical sign is intriguing in the context of the Braak-Del Tredici hypothesis, which implicates exposure to one or several pathogens in disease initiation. Moreover, the presence of αSyn in the mammalian olfactory epithelium and mucosa [16] renders the nasal passages a plausible site for a disease-initiating interaction between airborne pathogens, such as RNA viruses, and αSyn, a highly abundant brain protein that is prone to misfolding. Consistent with this possibility, animal studies have revealed an upregulation and accumulation of αSyn in response to intranasal SARS-CoV2 administration [17,18]. It is believed that the impact of the virus on αSyn metabolism may be imposed either via a direct interaction intracellularly or may be related to innate, immunomodulatory functions of extracellular αSyn [19]. With respect to the former, heparin-binding sites on the SARS-Cov-2 spike protein may act as facilitator to seed the aggregation of other heparin-binding proteins, including αSyn, as well as amyloid β-peptide and tau [20]. Further, SARS-CoV-2 proteins are capable of forming amyloid aggregates themselves [21], possibly providing a nidus for the aggregation of neurodegeneration-associated proteins. Alternatively, or in addition, αSyn has been demonstrated to function as an innate, anti-viral factor [16,22]. In theory, its upregulation in response to SARS-CoV2 infection could predispose it to aggregation, thereby putting COVID-19 patients at higher risk of developing PD-relevant changes [23]. Of note, few cases of parkinsonism associated with SARS-CoV-2 infection in humans have been documented since the beginning of the pandemic [24–26]. In contrast, a substantial number of patients with established PD experienced worsening of their symptoms during and after recovery from COVID-19 illness [27].

These lines of evidence implicate SARS-CoV-2 infection as a possible trigger for the initiation of PD-linked pathology. However, the scarcity of histopathological studies interrogating inflammatory and neurodegenerative changes and specifically probing for αSyn aggregation in the olfactory pathway following SARS-CoV-2 infection, represents a gap in our understanding of the relationship between the two disorders. In the present autopsy study, we employ an immunohistochemical approach to document the nature and extent of inflammation and of neurodegenerative changes within rostral olfactory structures, specifically the OB and OT, among a cohort of individuals that died of COVID-19-related complications versus those with neurodegenerative synucleinopathies, including PD, dementia with Lewy bodies (DLB), and multisystem atrophy (MSA). In addition, in light of reports that COVID-19 may predispose to Alzheimer disease (AD) [28], we have included participants with dementia as well as another tauopathy, progressive supranuclear palsy (PSP). We compared inflammatory and neurodegenerative changes in the OB and OT of these groups with a series of COVID-19-negative subjects without a neurodegenerative disease. Among the latter, we included as a further control a subset of participants with primary inflammatory disorders of the brain.

## Methods

### Study Subjects

Study subjects were recruited at two sites: The Ottawa Hospital in Ottawa, Ontario, Canada and the Kantonsspital St. Gallen, in St. Gallen, Switzerland. Ethics approval was obtained from the Ottawa Health Science Network Research Ethics Board (#20120963-01H) and the Kantonsspital St. Gallen (Ethikkommission Ostschweiz, Projekt-ID 2021-00678) respectively.

Characteristics of the study groups are summarized in **Table 1**. We examined tissue from 50 individuals: 16 COVID-19-negative controls, including 9 neurologically healthy participants (HCO) and 7 with inflammatory CNS disorders (referred to as neurological controls, NCO), 22 subjects who died from SARS-CoV-2 infection-related complications (referred to as COVID19+), 6 participants with Lewy Body disease (LBD), three with Alzheimer disease (AD), and three with other neurodegenerative diseases (OND; progressive supranuclear palsy (PSP), n=2; multisystem atrophy (MSA), n=1. The age of subjects ranged from 51 to 90 years. All COVID19+ patients were clinically free of a previously identified neurodegenerative illness based on medical chart review during their admission to the hospital, with the exception of one person, who demonstrated signs of parkinsonism, but without a formal diagnosis by a neurologist. Of note, olfactory function was not routinely measured in study participants; one COVID19+ patient reported the loss of sense of smell on admission. Details with respect to causes of death, clinical diagnoses and a summary of the sample inventory can be found in **Supplemental Table 1**.

**Table 1:** Demographics of study group.

|  | N | Age<br>(mean ± SD) | Age (yrs,<br>range) | Sex |  | PMI (hrs,<br>mean ± SD) |
| --- | --- | --- | --- | --- | --- | --- |
|  |  |  |  | M | F |  |
| Controls | 16 | 70.4 ± 8.2 | 55-84 | 7 | 9 | 68.3 ± 87.0 |
| Healthy Controls | 9 | 72.8 ± 7.5 | 59-80 | 4 | 5 | 52.1 ± 24.3 |
| Neurological Controls | 7 | 70.6 ± 10.5 | 55-84 | 3 | 4 | 89.1 ± 131.2 |
| COVID19+ | 22 | 74.8 ± 12.1 | 51-89 | 13 | 9 | 43.9 ± 25.2 |
| Lewy Body Disease | 6 | 78.7 ± 6.7 | 72-90 | 6 | 0 | 104.0 ± 111.2 |
| Alzheimer's Disease | 3 | 69.7 ± 15.3 | 52-78 | 1 | 2 | 56.0 ± 55.4 |
| Other Neurodegenerative Disease | 3 | 78.0 ± 3.6 | 74-79 | 3 | 0 | 112.0 ± 113.4 |

### Tissue processing

Tissues were removed and processed in the Departments of Pathology at The Ottawa Hospital and the Kantonsspital St. Gallen according to routine procedures for *post mortem* brain collection. Following their removal at autopsy, brain specimens were fixed typically for 10-14 days, rarely as long as 42 days, in 20% neutral buffered formalin. Following fixation, OBs and OT were dissected from the brain (unless they had already been separated at the time of removal of the rest of the brain) and embedded in paraffin.

### Immunohistochemistry

Sections were cut at 5 μm and mounted onto coated slides. Slides were deparaffinized in Citrisolv and rehydrated through a series of decreasing ethanol concentrations. Endogenous peroxidase activity was quenched with 0.3% hydrogen peroxide in methanol, followed by heat-induced antigen retrieval with citrate buffer, pH 6.0. For amyloid-β peptide staining, slides were also incubated in 98% formic acid for 5 minutes at room temperature. To reduce non-specific binding, sections were blocked in 10% goat serum in PBS-T (PBS + 0.1% Triton-X-100 + 0.05% Tween-20) for 30 minutes at room temperature. Sections were incubated overnight at 4 °C in primary antibodies diluted in 5% goat serum in PBS-T. Primary antibodies included those to: p-αSyn, pSyn#64 (Wako, cat# 015-25191, 1:500); LB509 (Biolegend, cat# 807702, 1:10,000); amyloid-β peptide, 6E10 (Biolegend cat# 803001, 1:1000); KiM1P (CD68; prepared in the Stadelmann lab; Radzun et al. 1991 [29]; 1:50); and SARS Nucleocapsid protein (Novus, cat# NB100-56576, 1:250). Biotinylated, secondary antibody anti-mouse or anti-rabbit IgG (H + L), made in goat (Vector Labs, BA-9200 or BA-1000) was diluted to 1:225, and sections were incubated for 1 hr at room temperature. The signal was amplified with VECTASTAIN® Elite® ABC HRP Kit (Vector Labs, PK-6100) for 1 hr at room temperature and visualized using 3’3-diaminobenzidine (Sigma, SIGMAFAST™ DAB, D4293). Samples were counterstained with Harris’ Modified Hematoxylin stain and dehydrated through a series of increasing ethanol concentration solutions and Citrisolv. Permount Mounting Medium (Fisher Scientific, SP15-100) was used for mounting, and developed slides were dried and scanned for visualization. The detailed IHC protocol can be found here: dx.doi.org/10.17504/protocols.io.kqdg3p7mql25/v1

Immunohistochemical staining for p-tau was performed in the Louise Pelletier Histopathology Core Facility in The Department of Pathology and Laboratory Medicine at The University of Ottawa using the Bond Polymer Refine Detection Kit (DS9800) with the Leica Bond™ system. Sections were deparaffinized and incubated using a 1:2500 dilution of mouse p-tau antibody (#MN1020, AT8 clone; ThermoFisher) for 15 minutes at room temperature and detected using an HRP conjugated compact polymer system. Slides were then stained using DAB as the chromogen, counterstained with Hematoxylin, mounted and cover slipped.

Image visualization across sites was performed using an *in-house* Omero Server software 5.6.6 [30] and immunohistochemistry figures were created using an *in-house* Omero Figure software v6.0.0.

Additional details of reagents can be found in **Supplemental Table 2.**

### Quantification

One section from each OB/OT was stained and analysed for each marker. In preliminary studies, it was evident that staining for neurodegenerative proteins was confined predominantly to the anterior olfactory nuclei (AON); thus, semi-quantitative scoring of immunoreactivity and intensity of inflammatory as well as neurodegenerative changes were focused on these nuclei, visually defined as groups of larger neurons lying on a neuropilic background (**Figure 1**). One section for each bulb was stained for CD68 reactivity as a surrogate marker for inflammation. For CD68 and neurodegenerative immunostaining, semi-quantitative analyses were performed using a scale from 0 to 5, corresponding to increasing densities of DAB-positive reactivity. **Supplemental Tables 3-7** indicate the number of AONs analyzed in each section, which tissue region was available for each case, as well as the score assigned for each marker used.

**Figure 1:**
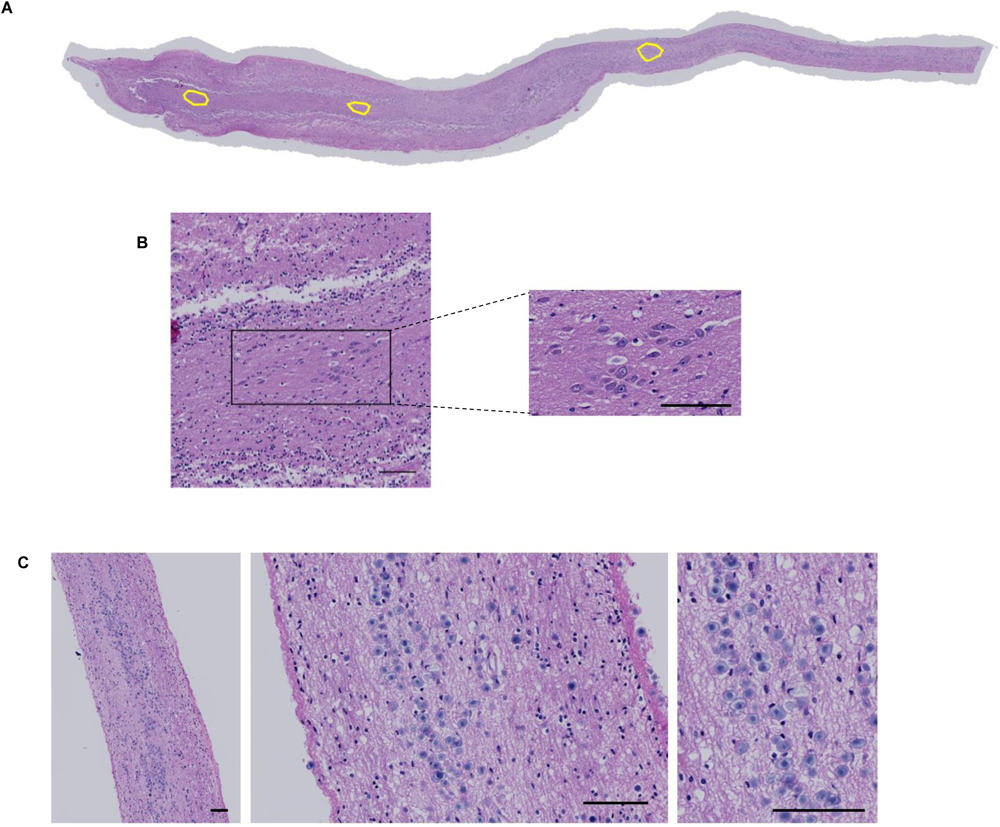
Overview of anatomical structures in the human olfactory bulb and tract. **A)** Hematoxylin and eosin (H&E)-stained section of a human olfactory bulb and tract [case #23] revealing three intrafascicular anterior olfactory nuclei (AON), as outlined in yellow, and in **B)** at higher magnification the most rostrally located AON is shown. **C)** H&E-stained section showing corpora amylacea (in blue) located in the subpial area of the human olfactory tract. Scale bars represent 100µM.

### Statistical analysis

Statistical analyses were performed using GraphPad Prism version 10 (GraphPad Software www.graphpad.com). Differences between groups were determined using the Kruskall-Wallis test followed by Dunn’s post-hoc analysis. Correlation between staining score and age was determined using Pearson’s correlation tool. For all statistical analyses, a cut-off for significance was set at 0.05. Data are displayed with p values represented as *p < 0.05, **p < 0.01, ***p < 0.001, and ****p < 0.0001.

## Results

Hematoxylin and eosin (HE)-stained sections of the OB and attached OT are shown in **Figures 1A** and **B**. These are representative of all tissue sections used in the analyses, and they indicate the location and appearance of the AON. As expected, in cases from older individuals, abundant corpora amylacea were often observed along white matter structures within the OT and were concentrated in the subpial neuropil (**Fig 1C**).

When we tested for SARS-CoV2 reactivity in sections of COVID19+ patients in our autopsy series using a specific antibody against its nucleocapsid protein, we did not detect any signal for the viral protein in any of the OB and OT sections analyzed under these conditions. In contrast, the same antibody readily detected the nucleocapsid protein in sustentacular cells of the olfactory epithelium in mice nasally inoculated with a mouse-adapted variant of SARS-CoV2 (not shown) [31].

The degree of tissue inflammation was assessed using anti-CD68 immunoreactivity for the detection of histiocytes and microglia. Positive CD68-immunoreactive cells were observed throughout the OB and OT (**Fig 2A**). To correlate inflammatory signals with neurodegenerative changes, we focused our semi-quantitative analysis of CD68 reactivity on the AON. As indicated in **Fig 2C**, all autopsy cases exhibited some degree of positive staining (a score of 1, or higher), regardless of disease status. For the purpose of comparing inflammatory changes, we divided the non-neurodegenerative controls into healthy controls (HCO) and those whose diagnosis at autopsy involved an inflammatory condition of the CNS (neurological controls; NCO). Because on average there was no significant difference in CD68 reactivity scores between HCO and NCO brains, these brains were grouped together as controls in subsequent analyses. Sections of AD brains generated the highest average inflammation score (4.7) in OB and OT, followed by the NCO group (average score, 4). COVID19+ cases had significantly lower CD68 staining scores than the NCO group (**Fig 2C**). Interestingly, inflammation scores based on CD68 staining correlated negatively (R=-0.4494) with age in COVID19+ patients (**Fig 2D**).

**Figure 2:**
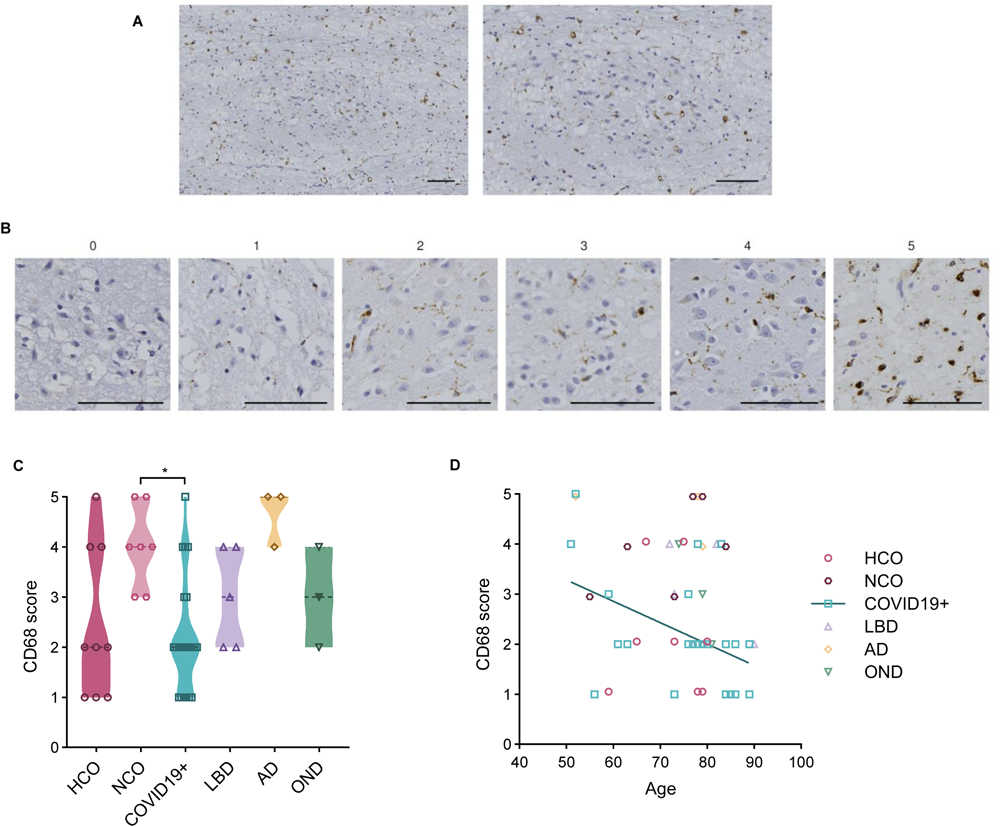
Anti-CD68 reactivity in the anterior olfactory nucleus. **A)** Example of immunohistochemical staining for CD68 in the human olfactory bulb [case #10]. **B)** Representative images of semi-quantitative scoring of staining, scale from 0 to 5, in the AON. **C)** Average staining score for each diagnostic group; filled triangle in the OND group indicates multiple system atrophy case. **D**) Scatter plot showing correlation between age and CD68 score for all groups. Scale bars represent 100µM. Significance was determined using Kruskal-Wallis test with Dunn’s post-hoc, where * indicates p≤0.05 (**C**) and Pearson’s correlation where r=-0.4494 (**D**). Straight line in D denotes correlation for COVID19+ cases. HCO denotes neurologically healthy controls; NCO, neurological controls; LBD, Lewy body disease; AD, Alzheimer disease; OND, other neurodegenerative diseases.

Phosphorylated αSyn (p-αSyn)-positive pathology was found to be restricted to the AON (**Fig 3A**). There, immunostaining was also scored on a semi-quantitative scale (as above) with representative images shown in **Fig 3B**. Not surprisingly, p-αSyn pathology was detected within the AON in all LBD cases (**Fig 3C**). Lewy neurites (LNs) and smaller, granular neuropil inclusions were found in several AON, but no definite, typical spherical Lewy bodies (LBs) within neuronal perikarya were observed. In the MSA case, abundant, pathognomonic glial cytoplasmic inclusions in oligodendrocytes were identified throughout the OB and OT (**Supp Fig 1**). Typical p-αSyn pathology was detected in four of 22 (18%) COVID19+ cases. Three subjects had no documented history of clinical signs of parkinsonism, whereas one was suspected to have parkinsonism (without a formal neurological diagnosis) (**Fig 3C**). In our series, anti-p-αSyn reactivity did not correlate with age (**Fig 3D**). Importantly, three of four anti-p-αSyn-positive COVID19+ subjects showed evidence of ‘incidental LBD’ in the form of LBs and LNs in the substantia nigra and dorsal motor nucleus of the vagus nerve. A fourth p-αSyn-positive COVID19+ case showed only a single reactive neurite in the OB, thus being assigned a score of 1. LBD cases were found to have a significantly higher average pathology score (4.3) than the control groups and COVID19+ cases (**Fig 3E**). One AD case with positive p-aSyn pathology staining was diagnosed with mixed pathology (AD plus LBD) when assessing the entire brain (**Fig 3C**). Immunohistochemical analyses using the anti-αSyn antibody, LB509, revealed similar results when compared to those using anti-p-aSyn (**Supp Fig 2**).

**Figure 3:**
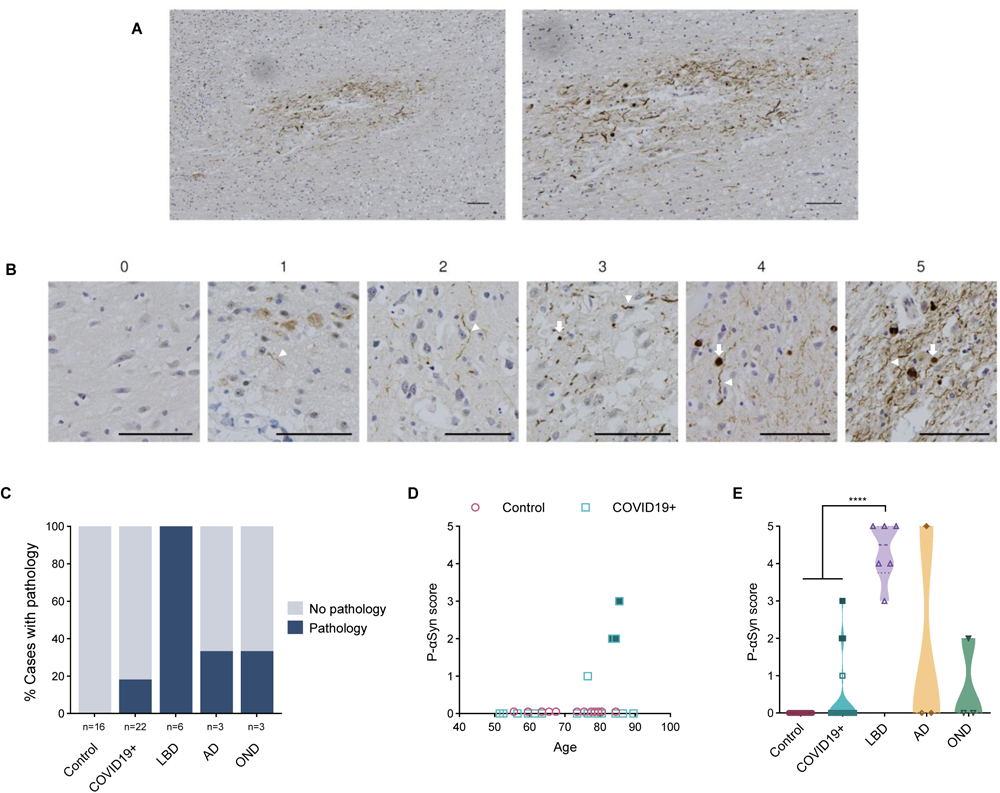
Anti-phosphorylated α-synuclein reactivity in the anterior olfactory nucleus. **A)** Example of immunohistochemical staining for p-αSyn in the human olfactory bulb, highlighting the AON from a subject with Parkinson disease dementia [case #39]. **B)** Representative images of semi-quantitative scoring of pathology, ranging from 0 to 5, in the AON. White arrows and arrowheads indicate Lewy inclusions and Lewy neurites, respectively. **C)** Percentage of cases in each group that have a pathology score of 1 or more. **D)** Correlation between age and p-αSyn pathology scores in the control group (HCO and NCO) and COVID19+ cases. **E)** Distribution of pathology scores for each group. Filled blue squares in D and E indicate COVID19+ cases suspected of having incidental LBD at autopsy; filled dark yellow diamond indicates AD case diagnosed with mixed pathology at autopsy; filled green triangle indicates MSA case. Scale bars represent 100µM. Significance was determined using Kruskal-Wallis test with Dunn’s post-hoc (**E**), where **** indicates p≤0.0001. Abbreviations for disease groups as in Figure 1.

Phosphorylated tau (p-tau) inclusions were detected by the AT8 antibody in 90% of the cases and were confined to the AON (**Fig 4A**). These tau-immunoreactive aggregates included dystrophic neurites as well as neurofibrillary tangles. Representative images corresponding to the semi-quantitative scale used for scoring (as above) are shown in **Fig 4B**. There was no correlation between the severity of p-tau pathology and the age of neurologically healthy cases (**Fig 4D**). AD cases had the highest average pathology score (5.0) among all groups, which was significantly higher than reactivity seen in the control and COVID19+ groups. (**Fig 4E**).

**Figure 4:**
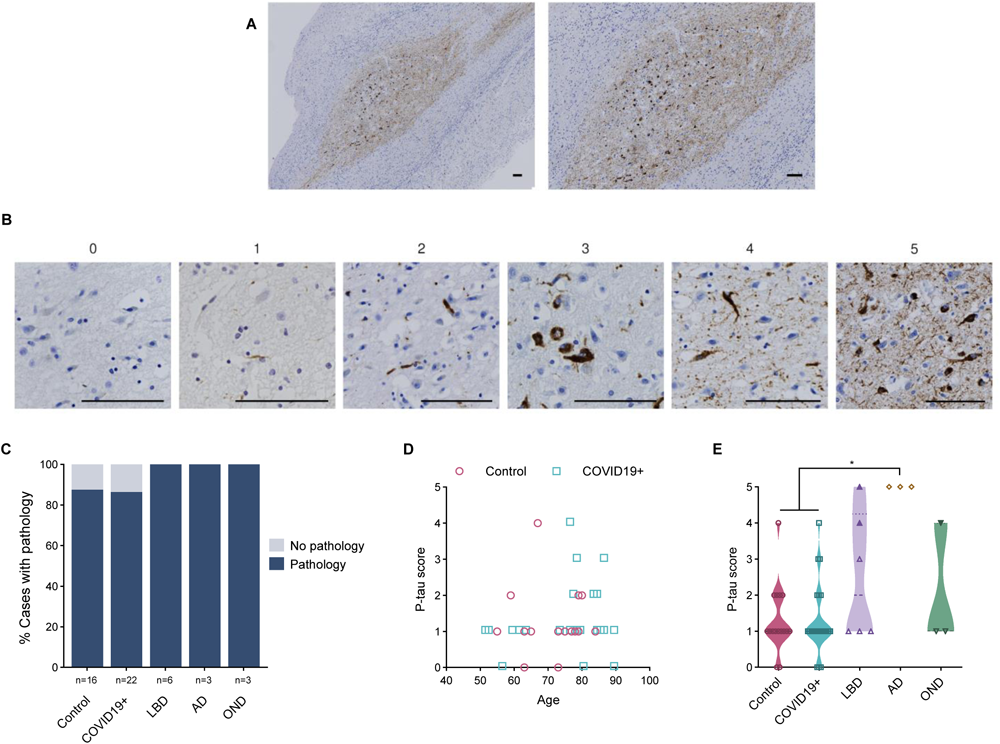
Anti-phosphorylated tau reactivity in the anterior olfactory nucleus. **A)** Example of immunohistochemical staining for p-tau in the human olfactory bulb, concentrated in the AON, from a subject with Alzheimer’s disease [case #47]. **B)** Representative images of semi-quantitative scoring of pathology, from 0 to 5, in the AON. **C)** Percentage of cases in each group that have a pathology score of 1 or more. **D)** Correlation between age and p-tau scores for the control (HCO and NCO) and COVID19+ groups. **E)** Distribution of tau pathology score for each group; filled purple triangles indicate LBD cases diagnosed with mixed pathology at autopsy. Filled green triangle indicates MSA case. Scale bar represents 100µM. Significance was determined using Kruskal-Wallis test with Dunn’s post-hoc, where * indicates p≤0.05. Abbreviations for disease groups as in Figure 1.

When detected, amyloid-β peptide-specific pathology was found to be restricted to the AON (**Fig 5A**), with representative images shown in **Fig 5B**. As expected, all cases in the AD group were found to have amyloid-β pathology. A small percentage of control and COVID19+ cases were found to have amyloid-β plaques in the AON, while other neurodegenerative cases did not show any detectable amyloid-β-positive pathology in their OB or OT (**Fig 5C**). There was no correlation between the degree of amyloid-β peptide burden and age in control or COVID19+ cases (**Fig 5D**). One NCO case (grouped here together with HCO) with amyloid-β-related angiitis generated an amyloid-β score of 3. The COVID19+ brain that contained the highest amyloid-β peptide load showed AD neuropathologic changes on *post mortem* examination of other brain regions despite the absence of a documented clinical history of dementia. When looked at as a group, AD cases had a significantly higher average score for amyloid-β deposition (4.7), as expected, than corresponding tissues from controls, COVID19+ and OND subjects (**Fig 5E**).

**Figure 5:**
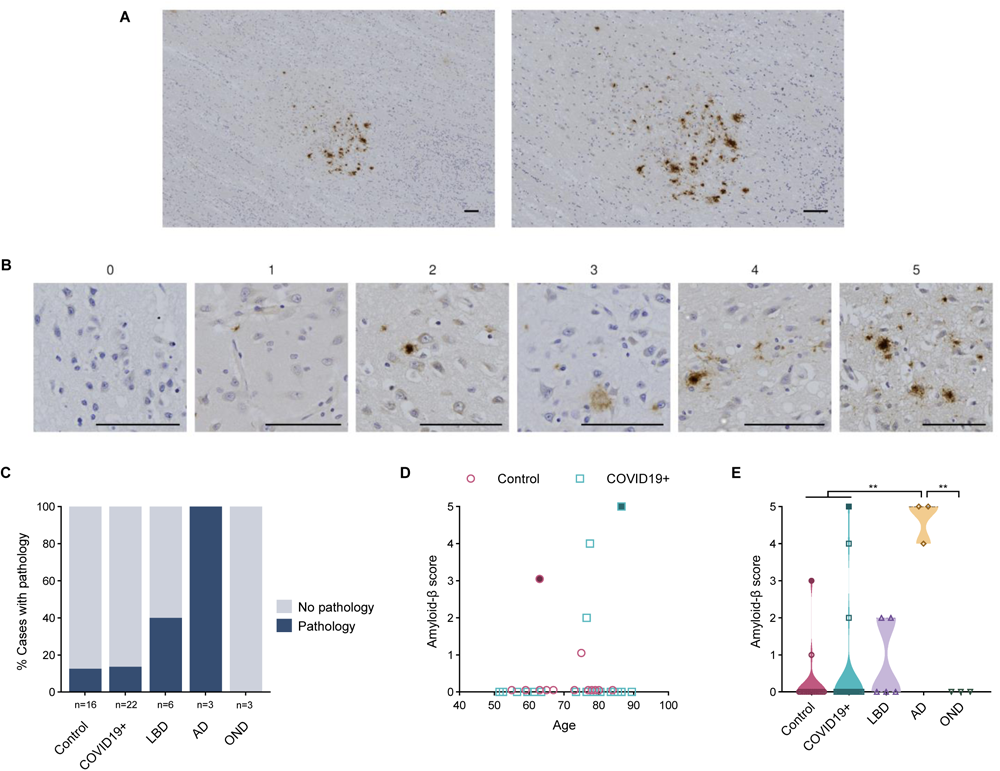
Anti-amyloid-β peptide reactivity in the anterior olfactory nucleus. **A)** Example of immunohistochemical staining for amyloid-β peptide in human olfactory bulb, concentrated in the AON, from a subject with Alzheimer’s disease [case #46]. **B)** Representative images of semi-quantitative scoring of pathology, from 0 to 5, in the AON. **C)** Percentage of cases in each group that have a pathology score of 1 or more. **D)** Correlation between age and amyloid-β peptide scores in the control (HCO and NCO) and COVID19+ groups. Subject highlighted in blue is suspected to have Alzheimer’s disease (AD) based on neuropathological findings in the temporal lobes. **E)** Distribution of pathology score for each group. Filled maroon circle indicates a control subject with amyloid-β-related angiitis; filled blue square indicates COVID19+ case suspected of having AD. Scale bar represents 100µM. Significance was determined using Kruskal-Wallis test with Dunn’s post-hoc, where ** indicates p≤0.01. Abbreviations for disease groups as in Figure 1.

Lastly, we sought to correlate staining for neurodegenerative proteins with the degree of CD68-immunoreactivity (**Fig 6A-C**). When comparing the cases with a pathology score of 1 or above to those with no pathology (score 0), p-tau cases with a score of 5 had significantly higher CD68 reactivity (**Fig 6B**). Individual trends but no significant differences were seen when correlating anti-CD68 immunoreactivity scores with the degree of synucleinopathy or amyloid-β peptide amyloidosis (**Fig 6A,C**).

**Figure 6:**
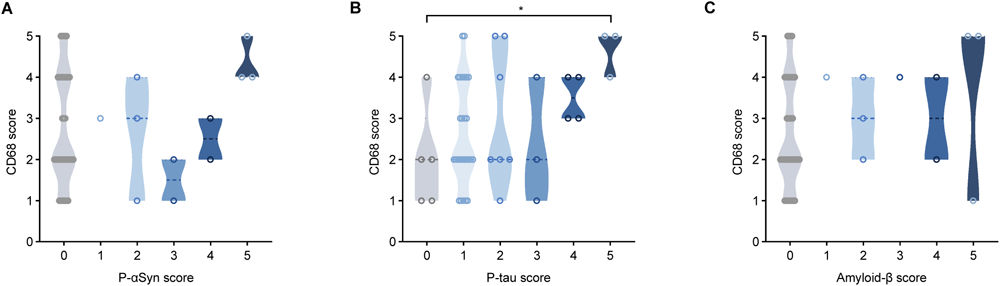
Correlation studies between inflammation and neurodegeneration-linked pathology in the anterior olfactory nucleus. Violin plots showing relationships between scores for anti-CD68 immunoreactivity and three neurodegeneration-linked proteins in the AON, namely phosphorylated α-synuclein (**A**), phosphorylated tau (**B**) and amyloid-β peptide (**C**). Significance was determined using Kruskal-Wallis test with Dunn’s post-hoc, using a score of 0 (indicating absent pathology) for comparison, where * indicates p≤0.05.

Individual scores for the immunostaining with each antibody for each of the 50 cases are shown in **Table 2**, with additional details summarized in **Supplemental Tables 3**-**7**.

**Table 2:** Heat map of all staining scores for each case in this study. Characteristics listed include age (in years); sex (F, female; M, male); diagnostic group (HCO, healthy control; NCO, neurological control; LBD, Lewy Body Disease’ AD, Alzheimer’s disease; OND, other neurological disease); and pathology scores in the AON (anterior olfactory nucleus) for each marker, from 0 to 5.

| Case ID | Age | Sex | Group | Staining score in AON (from 0-5) |  |  |  |
| --- | --- | --- | --- | --- | --- | --- | --- |
| | | | | P-αSyn | P-tau | Amyloid-β | CD68 |
| 1 | 55 | F | NCO | 0 | 1 | 0 | 3 |
| 2 | 59 | M | HCO | 0 | 2 | 0 | 1 |
| 3 | 63 | M | NCO | 0 | 0 | 0 | 4 |
| 4 | 63 | M | NCO | 0 | 1 | 3 | 4 |
| 5 | 65 | M | HCO | 0 | 1 | 0 | 2 |
| 6 | 67 | F | HCO | 0 | 4 | 0 | 4 |
| 7 | 73 | F | HCO | 0 | 0 | 0 | 2 |
| 8 | 73 | F | NCO | 0 | 1 | 0 | 3 |
| 9 | 75 | F | HCO | 0 | 1 | 1 | 4 |
| 10 | 77 | M | NCO | 0 | 1 | 0 | 5 |
| 11 | 78 | M | HCO | 0 | 1 | 0 | 1 |
| 12 | 79 | F | HCO | 0 | 1 | 0 | 1 |
| 13 | 79 | F | HCO | 0 | 2 | 0 | 5 |
| 14 | 79 | F | NCO | 0 | 2 | 0 | 5 |
| 15 | 80 | M | HCO | 0 | 2 | 0 | 2 |
| 16 | 84 | F | NCO | 0 | 1 | 0 | 4 |
| 17 | 51 | M | COVID19+ | 0 | 1 | 0 | 4 |
| 18 | 52 | M | COVID19+ | 0 | 1 | 0 | 5 |
| 19 | 56 | M | COVID19+ | 0 | 0 | 0 | 1 |
| 20 | 59 | M | COVID19+ | 0 | 1 | 0 | 3 |
| 21 | 61 | F | COVID19+ | 0 | 1 | 0 | 2 |
| 22 | 63 | F | COVID19+ | 0 | 1 | 0 | 2 |
| 23 | 73 | F | COVID19+ | 0 | 1 | 0 | 1 |
| 24 | 76 | M | COVID19+ | 1 | 4 | 2 | 3 |
| 25 | 76 | F | COVID19+ | 0 | 1 | 0 | 2 |
| 26 | 77 | F | COVID19+ | 0 | 2 | 4 | 2 |
| 27 | 78 | M | COVID19+ | 0 | 3 | 0 | 2 |
| 28 | 78 | F | COVID19+ | 0 | 1 | 0 | 4 |
| 29 | 79 | M | COVID19+ | 0 | 1 | 0 | 2 |
| 30 | 80 | F | COVID19+ | 0 | 0 | 0 | 2 |
| 31 | 83 | F | COVID19+ | 2 | 2 | 0 | 4 |
| 32 | 84 | M | COVID19+ | 2 | 1 | 0 | 1 |
| 33 | 84 | M | COVID19+ | 0 | 2 | 0 | 2 |
| 34 | 85 | M | COVID19+ | 3 | 1 | 0 | 1 |
| 35 | 86 | M | COVID19+ | 0 | 3 | 5 | 1 |
| 36 | 86 | F | COVID19+ | 0 | 1 | 0 | 2 |
| 37 | 89 | M | COVID19+ | 0 | 1 | 0 | 2 |
| 38 | 89 | M | COVID19+ | 0 | 0 | 0 | 1 |
| 39 | 72 | M | LBD | 5 | 4 | 2 | 4 |
| 40 | 73 | M | LBD | 4 | 1 | 0 | 3 |
| 41 | 77 | M | LBD | 4 | 1 | 0 | 2 |
| 42 | 79 | M | LBD | 5 | 5 | - | - |
| 43 | 82 | M | LBD | 5 | 3 | 0 | 4 |
| 44 | 90 | M | LBD | 3 | 1 | 2 | 2 |
| 45 | 52 | F | AD | 5 | 5 | 5 | 5 |
| 46 | 78 | F | AD | 0 | 5 | 5 | 5 |
| 47 | 79 | M | AD | 0 | 5 | 4 | 4 |
| 48 | 74 | M | OND | 0 | 1 | 0 | 4 |
| 49 | 79 | M | OND | 2 | 4 | 0 | 3 |
| 50 | 81 | M | OND | 0 | 1 | 0 | 2 |

## Discussion

Previous authors have pointed at a possible association between SARS-CoV-2 infection and neurodegeneration, as identified by a rise in the incidence of cognitive decline and parkinsonism in survivors [28,32–34]. A recent imaging study of COVID19+ subjects, including those with mild disease, revealed reductions in grey matter thickness in olfaction-related structures as well as other regions functionally connected to the olfactory cortex [15]. In contrast, other investigators [35,36] reported increases in gray matter volumes in distinct anatomical regions. These divergent results likely reflect substantial complexity in the relationship between pathogenic SARS-CoV-2 variants, genetic susceptibility of the host, mechanisms of disease, and the period that has passed between infection and assessment of outcomes.

Here, we did not find any evidence for a relationship between the history of a recent COVID-19-related illness, which had warranted hospital admission and was ultimately fatal, and features of αSyn pathology in the OB or AON, using immunohistochemical staining with two well-characterized, specific antibodies as a readout. Although we detected αSyn aggregates within the AON in four of 22 (18%) of the neurologically intact COVID19+ group, three of these subjects showed typical Lewy pathology in the brainstem, with only one of those having had suspected parkinsonism as per the medical record. It is generally thought that these hallmark findings of PD are generated over a decade (or more) [37], whereas the course of COVID-19 illness that lead to death in our study subjects was on average two weeks following hospitalization (**Supplemental Table 1**). Therefore, we speculate that the observed pathology in these cases reflected pre-existing LBD that occurred independently of SARS-CoV-2 infection. In this respect, our results are compatible with those of Blanco-Palmero and co-workers, who observed no significant difference in total αSyn levels in the cerebrospinal fluid or serum among COVID-19 patients versus HCO subjects [38].

However, our findings do not exclude an effect of the viral infection on subtle αSyn misprocessing in the olfactory system, as it is conceivable that there was insufficient time for immunohistochemically detectable, fibrillar αSyn aggregates to form. An implication of this caveat is that in the future more sensitive approaches for the detection of soluble oligomers, of post-translationally modified conformers, of in vitro seeding-competent species or of alternatively spliced mRNA transcripts could reveal such an association, including in the early stages after COVID-19 infection.

Previous *post mortem* analyses of COVID-19 patients had revealed local inflammation associated with axonal pathology and microvascular changes in the OB and OT [14], as well as the finding of increased immune activation and microglial nodules, as reported for example by Schwabenland et al. [39]. In our study, the absence of significant inflammation in COVID19+ subjects relative to controls, as judged by using CD68 immunoreactivity as a readout, is consistent with findings by Matschke et al. [40]. In our case series and akin to the evidence regarding αSyn aggregation, this relative paucity of inflammation within the OB and OT could also be related to the relatively short duration of illness, its transient nature, or lack of actual tissue invasion by virions. Conversely, one could speculate that less visible inflammation may reflect a relatively deficient immune response in older subjects, rendering them more susceptible to COVID-19 infection in general. Along these lines, we found that CD68 scores in the rostral olfactory circuitry were negatively associated with age in our COVID19+ cases, possibly reflecting a decline in anti-viral responses. Studies are underway to examine the degree of inflammation in the piriform cortex, to which the olfactory tracts project.

One shortcoming of our study is that no quantification of olfactory function had been performed in any of the participants that subsequently died within the hospital, in large part due to the severity of respiratory distress that warranted intubation. In a parallel effort, we are currently analyzing serial sections of the intact nasal cavity, including olfactory and respiratory epithelia, from all 50 study participants, where we will revisit the rate of detection of SARS-CoV-2 nucleocapsid proteins. When probing for the presence of viral proteins in OB and OT sections from three, PCR-confirmed COVID19+ cases in our series, we detected no specific evidence of a viral infection.

Although we were unable to procure evidence for a direct relationship between COVID-19 infection and olfactory αSyn aggregation, our study revealed interesting findings of relevance to olfactory dysfunction in neurodegeneration and perhaps to ageing as well. For example, our demonstration that the overall severity of αSyn as well as amyloid-β and tau pathology of the OB and AON was highest in the LBD and AD cases, respectively, is consistent with the frequent association of hyposmia with these diseases. Conversely, the relative paucity of αSyn and tau pathology within AON neurons from our MSA and PSP subjects, respectively, correlates with the absence of a loss in sense of smell as a cardinal clinical manifestation in these diseases [41].

The existence of tau and αSyn co-pathology in almost all of the synucleinopathy cases in our series suggests that the molecular mechanisms which underlie the frequent co-existence of these changes in the brains of AD and LBD patients are also operative in the intracranial olfactory circuitry.

Our rationale for including subjects with AD derived from evidence that SARS-CoV-2 infection may facilitate cognitive decline [28] and may disrupt both amyloid-β and tau homeostasis, leading to tau hyperphosphorylation, as suggested in part by elevated plasma concentrations of both total and phosphorylated tau in living subjects [42–45]. We detected no strong evidence yet to link COVID-19 infection to β-amyloidosis in the rostral olfactory circuitry. However, three COVID19+ cases (13.6%) compared to a single control individual (6.25%) were found to have amyloid-β pathology in the OB and OT. Although the numbers of cases in each diagnostic group are too small to draw definitive conclusions, this result is intriguing considering studies that showed SARS-CoV-2, herpetiform DNA viruses and bacteria can alter amyloid-β homeostasis and modify neuropathological outcomes [42,46].

We detected no evidence for a relationship between COVID-19 illness and p-tau deposition in the OB or AON at autopsy. Consistent with recent findings by Tremblay and colleagues [47], 90% of our cohort, including control participants, displayed at least some degree of tau pathology. This is perhaps not surprising given that the process of tau hyperphosphorylation and aggregation is not confined to primary neurodegenerative diseases. Indeed, changes in tau metabolism are increasingly recognized as sequelae of diverse insults to the brain, including seizures, trauma, viral infections and autoimmune disease [48,49], and as a part of physiological ageing [47]. In this context, the prevalence of tau pathology in the OB and AON across our entire cohort may represent a morphological surrogate for a variety of insults, including pathogen-induced inflammation, sustained by individuals throughout their lifetime. Perhaps consistent with this, tauopathy cases in our case series showed the highest inflammation (anti-CD68 reactivity-based) scores (**Fig 6B**). However, assuming that p-tau aggregation and inflammation may be related, it is not possible to ascribe temporal precedence to one or the other due to the cross-sectional nature of our study.

In conclusion, we found no evidence that a subacute, fatal SARS-CoV-2 infection induces αSyn aggregation in the OB or AON of human subjects under the microscopic conditions examined. However, our study has limitations and further investigations addressing these are required prior to excluding a role for this RNA virus in triggering neurodegeneration within the olfactory system. The most critical of these is the discrepancy between the duration of illness, which in our cohort was short, and the time required for neurodegenerative changes to be detectable by traditional immunohistochemistry. Related to this, the subjects in our cohort were all above 50 years of age, thereby complicating the attribution of neurodegenerative changes to SARS-CoV-2 alone versus those related to ageing, including those clinically not yet detected. Future studies should include more subjects, including younger individuals, those with a longer duration of viral illness and those with neurological signs as a result of their infectious illness. We will supplement histological analyses with more sensitive, biochemically based techniques and quantification of transcriptional changes to detect pathologically relevant structural and molecular alterations in the proteins of interest. In our prospectively collected cohort, there were no female subjects in our LBD and OND groups, precluding analysis of sex-related differences in these subsets. Further, because this post mortem study was cross-sectional, it precluded analyses of any temporal relationship(s) among the changes observed.

To address any long-term sequelae versus short-term effects of RNA virus infections of the upper respiratory tract on brain health, such as pertaining to αSyn, tau and amyloid-β peptide homeostasis, we have recently begun to conduct experiments in mice, including those that carry PD-linked allelic variants [16,50]. These ongoing studies in animal models include the inoculation of the nasal cavity with a mouse-adapted strain of SARS-CoV-2 [31] to monitor its potential aggregation effects on neurodegeneration-linked proteins. As mentioned, our investigation here was confined to the OB and AON. Autopsy studies interrogating the impact of SARS-CoV-2 infection in humans on αSyn metabolism (and inflammation) at a more rostral site in the olfactory epithelium as well as more proximally in the piriform cortex are also currently ongoing. We anticipate that these will add to an ultimately anatomically more complete assessment of this potentially important environment-brain interaction.

## Supplementary Information

### Acknowledgments

This research was funded by Aligning Science Across Parkinson’s [Grant ID: ASAP-020625] through the Michael J. Fox Foundation for Parkinson’s Research (MJFF) and by the Parkinson Research Consortium Ottawa. For the purpose of open access, the author has applied a CC BY public copyright license to all Author Accepted Manuscripts arising from this submission.

We gratefully acknowledge histology, staining, and imaging services provided by the Louise Pelletier HCF (RRID: SCR_021737) at the University of Ottawa and the support by Dr. Jay Maxwell and staff at The Ottawa Hospital Department of Pathology.

### Author Contributions

NAL, JJT, JF, CSN, aSCENT-PD Investigators, JMW and MGS conceptualized the study; NAL, carried out the histological assessments; NAL, JJT, JMW and MGS analyzed data; AKJ, JF, CSN, LF, OH, WJ, JP, JMW collected autopsy material, NAL, JMW, JJT, MGS wrote the manuscript and all authors provided feedback on manuscript.

---

**Supplemental Figure 1:**
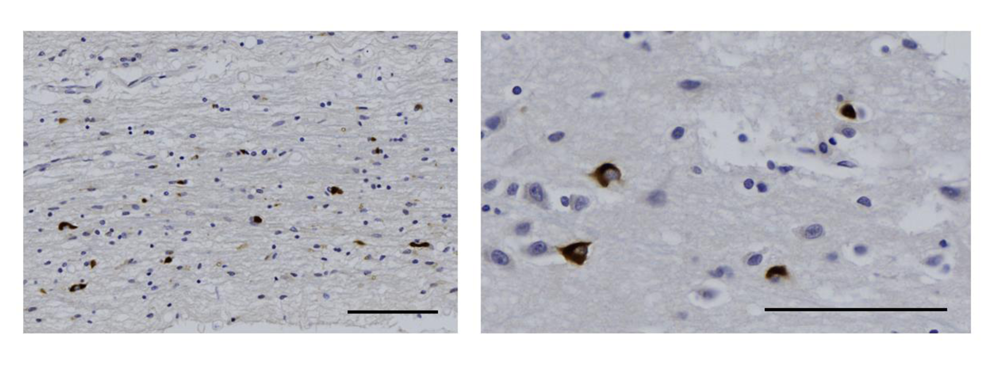
Example of phosphorylated α-synuclein pathology shown at low and high magnifications depicting glial cytoplasmic inclusions in oligodendrocytes of the anterior olfactory nucleus in the OB of an individual with multiple system atrophy (type P). Scale bars represent 100µM.

**Supplemental Figure 2:**
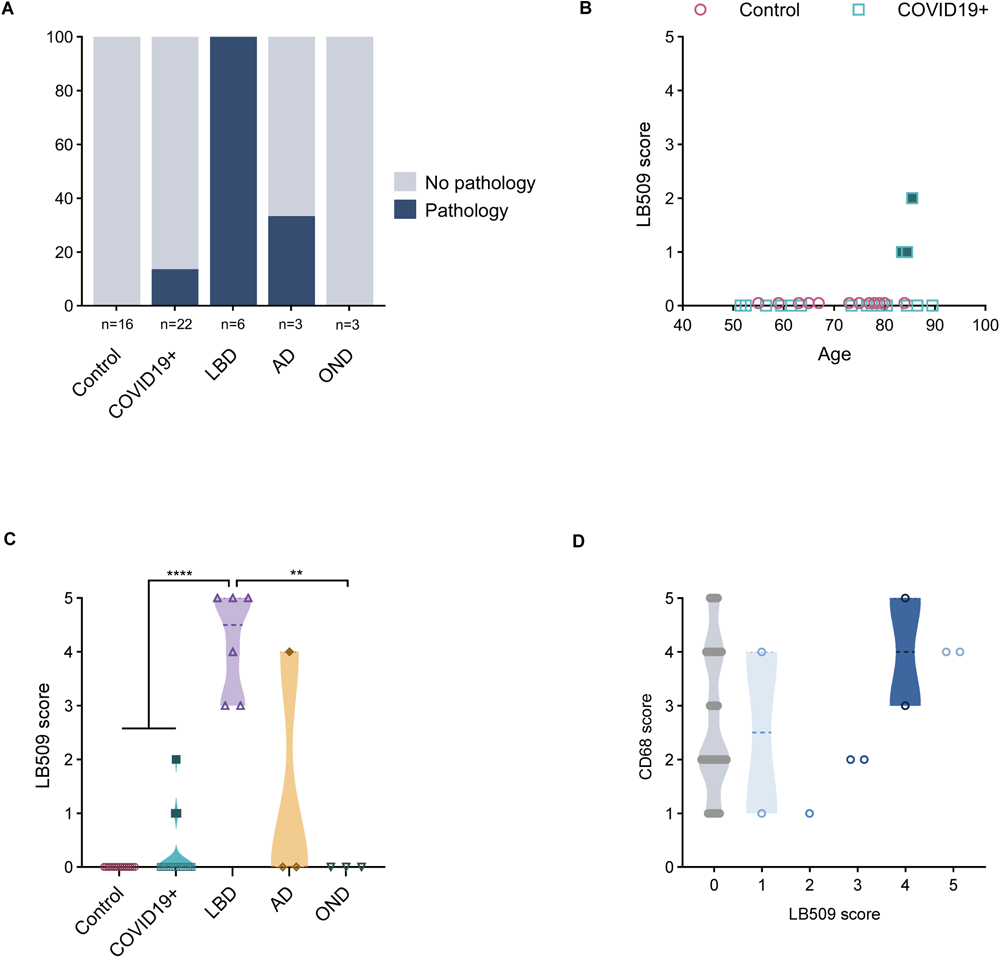
LB509-mediated α-synuclein reactivity in the anterior olfactory nucleus. **A)** Percentage of cases in each group that have a pathology score of 1 or more. **B**) Correlation between age and LB509 α-synuclein pathology scores in the control (HCO and NCO) and COVID19+ groups. **C**) Distribution of LB509 pathology scores for each group. Filled blue squares are subjects suspected of having incidental LBD, filled dark yellow diamond reflects a subject with mixed pathology. **D**) Relationship between anti-CD68 score and LB509 α-synuclein pathology score in the AON of all groups. Significance was determined using Kruskal-Wallis test with Dunn’s post-hoc, where ** denotes p≤0.01 and **** denotes p≤0.0001. Abbreviations for disease groups as in Figure 1.

**Supplemental Table 1:**
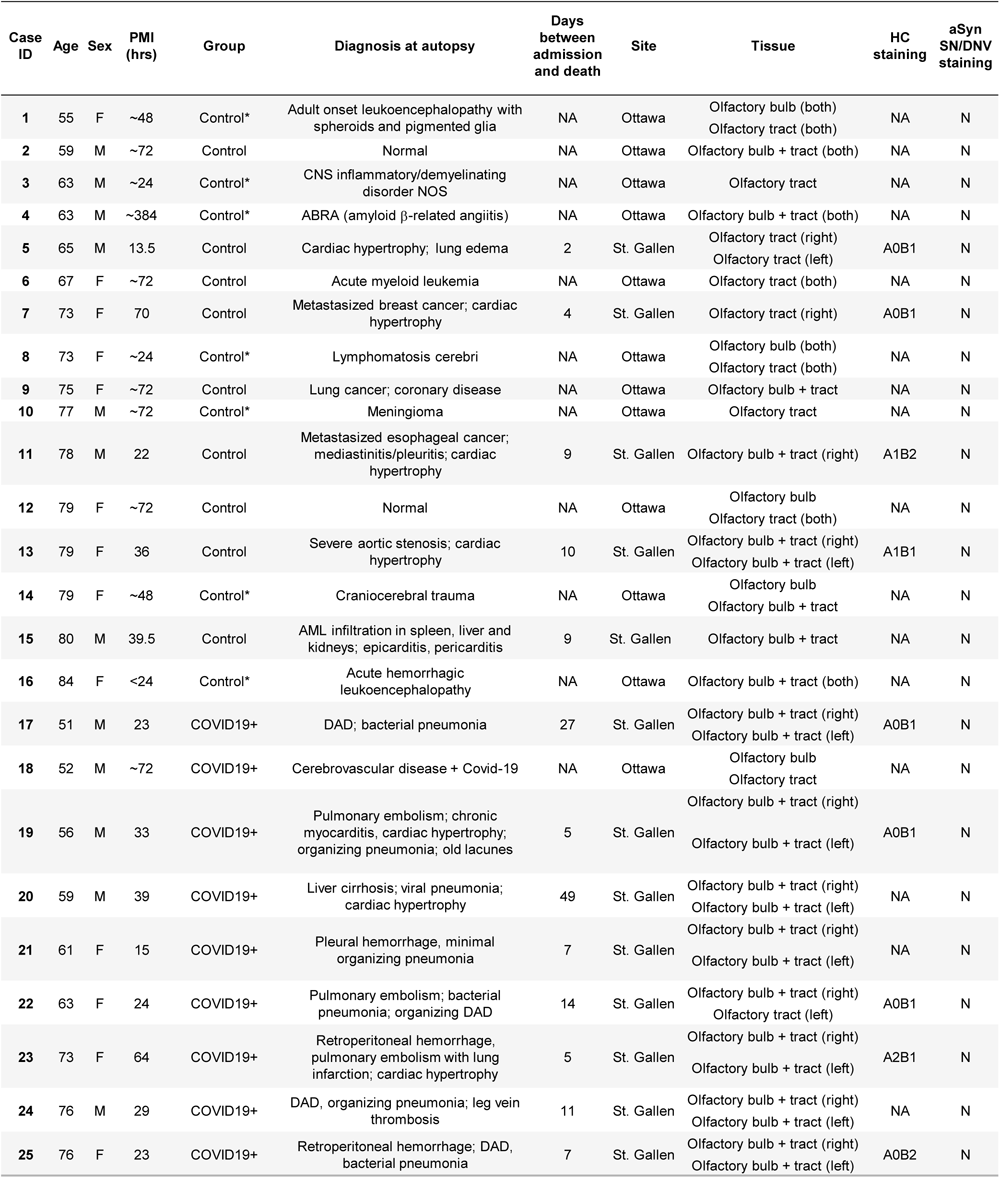
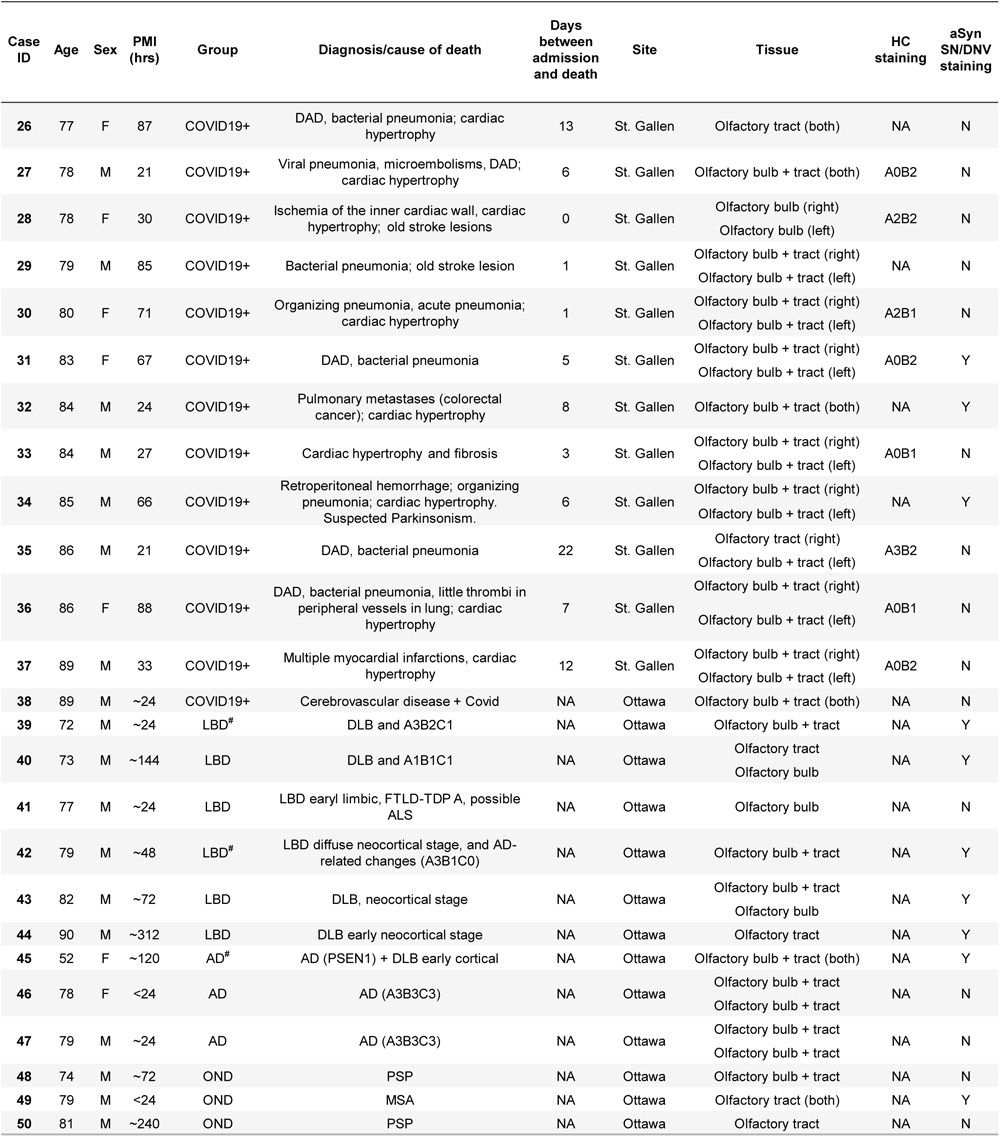
Detailed characteristics of study group. Characteristics listed include age (in years); sex (F, female; M, male); *post mortem* interval (PMI; in hours); diagnostic group (HCO, healthy control; NCO, neurological control; LBD, Lewy Body Disease’ AD, Alzheimer’s disease; OND, other neurological disease); cause of death or diagnosis at autopsy; number of days in hospital, NA, not applicable; site of tissue collection; tissue analyzed; hippocampal (HC) staining; aSynuclein staining in the substantia nigra (SN) and dorsal motor nucleus of the vagus (DNV).*Indicate cases with an inflammatory condition. ^#^Indicate cases with mixed pathology.

**Supplemental Table 2:** Resources and reagents used in study.

| Reagent or Resource | Source | Identifier |
| --- | --- | --- |
| **Antibodies** |  |  |
| Anti-phosphorylated αSynuclein | FUJIFILM Wako Shibayagi | Cat# 015-25191<br>RRID:AB_2537218 |
| Phospho-Tau (Ser202, Thr205) Monoclonal Antibody (AT8) | Thermo Fisher Scientific | Cat# MN1020<br>RRID:AB_223647 |
| KiM1P | Non-commercial. Gift from J. Franz | As described by Radzun et al. 1991 |
| Purified (azide-free) anti-beta-Amyloid, 1-16 | Biolegend | Cat# 803001<br>RRID:AB_2564653 |
| LB509 [Purified (azide-free) anti-alpha-Synuclein, 115-121] | Biolegend | Cat# 807702<br>RRID:AB_2564725 |
| SARS Nucleocapsid protein | Novus Biologicals | Cat# NB100-56576<br>RRID: AB_838838 |
| Anti-Mouse IgG (H+L), made in goat | Vector Laboratories | Cat# BA-9200<br>RRID:AB_2336171 |
| Anti-Rabbit IgG (H+L), made in goat | Vector Laboratories | Cat# BA-1000<br>RRID:AB_2313606 |
| **Reagent** |  |  |
| VECTASTAIN Elite ABC-Peroxidase Kit | Vector Laboratories | Cat# PK-6100<br>RRID:AB_2336819 |
| SIGMAFAST™ 3,3'-Diaminobenzidine tablets | Millipore Sigma | Cat# D4293 |
| PermOUNT Mounting Medium | Fisher Scientific | Cat# SP15-100 |

**Supplemental Table 3:**
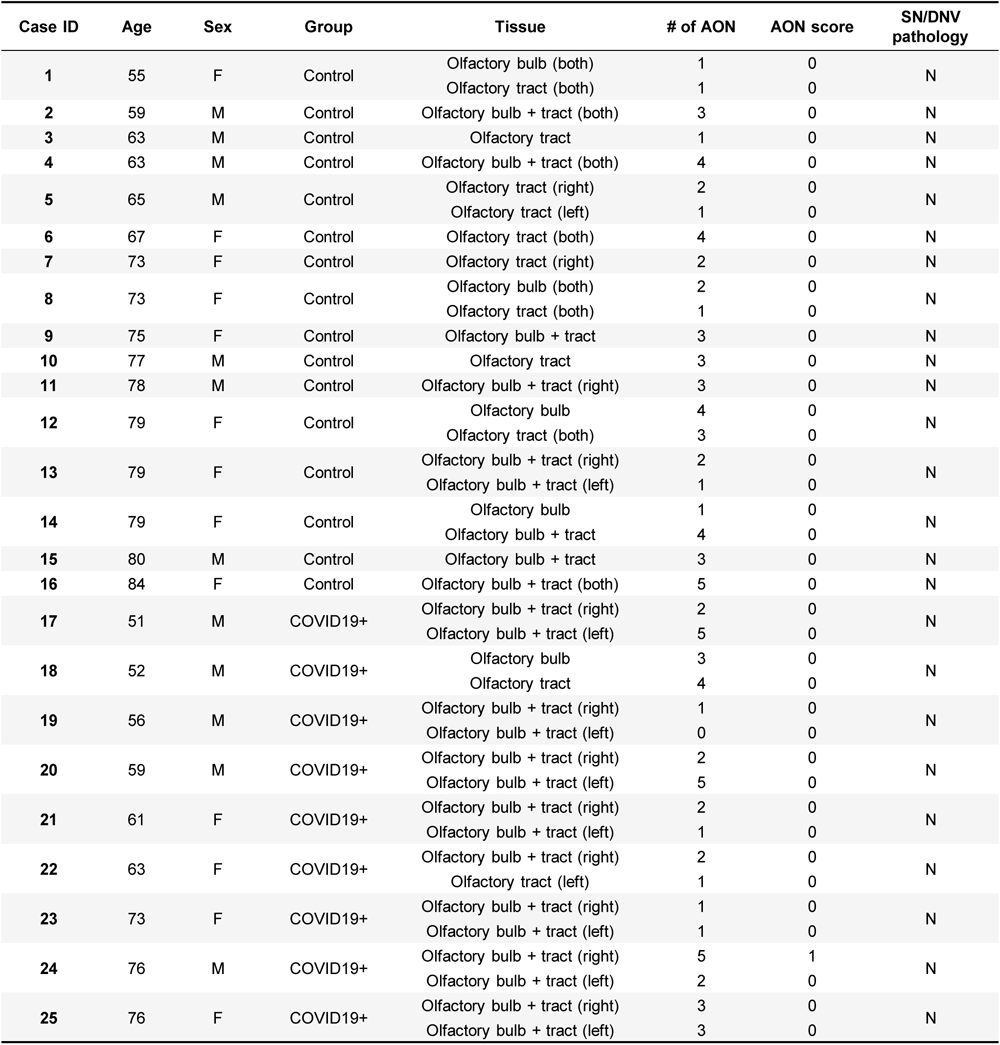
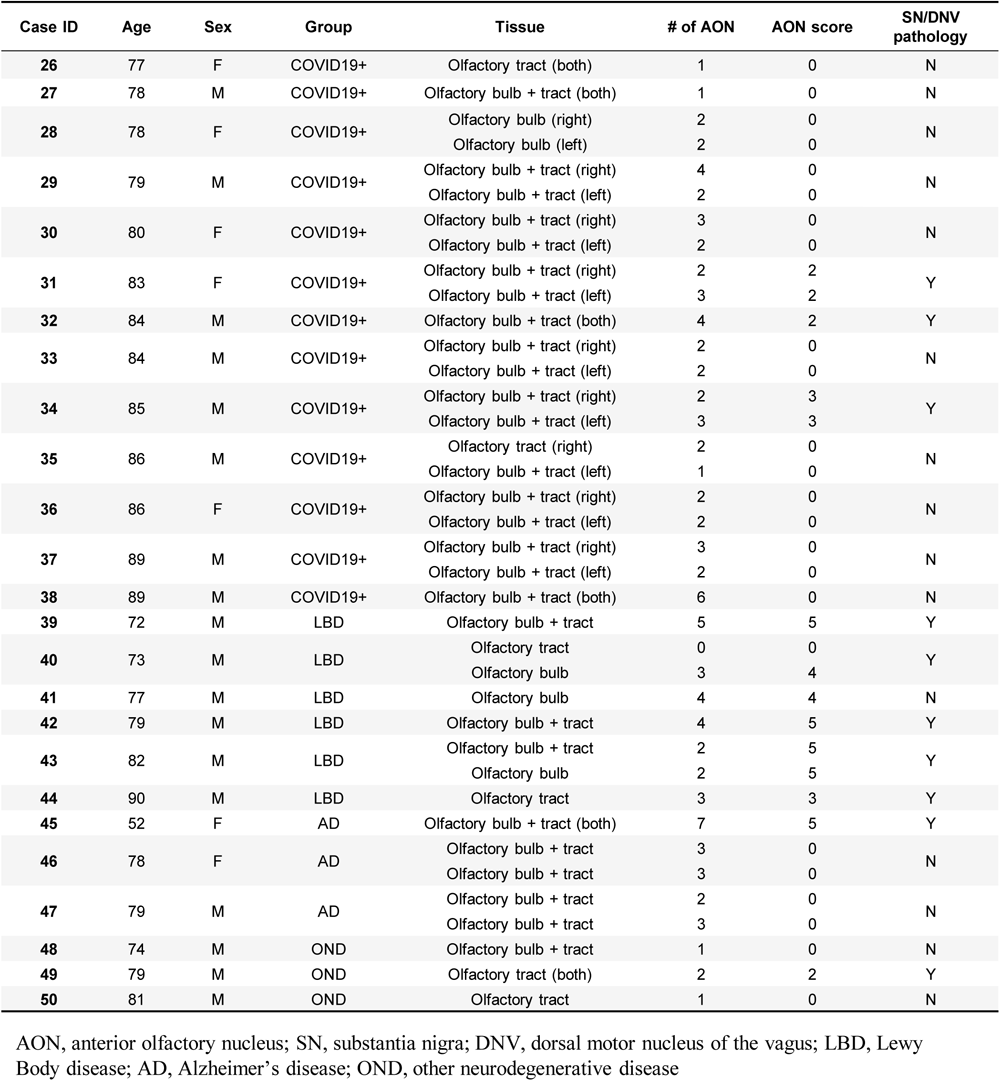
Phosphorylated α-Synuclein staining.

| Case ID | Age | Sex | Group | Tissue | # of AON | AON score | SN/DNV pathology |
| --- | --- | --- | --- | --- | --- | --- | --- |
| 1 | 55 | F | Control | Olfactory bulb (both) | 1 | 0 | N |
|  |  |  |  | Olfactory tract (both) | 1 | 0 |  |
| 2 | 59 | M | Control | Olfactory bulb + tract (both) | 3 | 0 | N |
| 3 | 63 | M | Control | Olfactory tract | 1 | 0 | N |
| 4 | 63 | M | Control | Olfactory bulb + tract (both) | 4 | 0 | N |
| 5 | 65 | M | Control | Olfactory tract (right) | 2 | 0 | N |
|  |  |  |  | Olfactory tract (left) | 1 | 0 |  |
| 6 | 67 | F | Control | Olfactory tract (both) | 4 | 0 | N |
| 7 | 73 | F | Control | Olfactory tract (right) | 2 | 0 | N |
| 8 | 73 | F | Control | Olfactory bulb (both) | 2 | 0 | N |
|  |  |  |  | Olfactory tract (both) | 1 | 0 |  |
| 9 | 75 | F | Control | Olfactory bulb + tract | 3 | 0 | N |
| 10 | 77 | M | Control | Olfactory tract | 3 | 0 | N |
| 11 | 78 | M | Control | Olfactory bulb + tract (right) | 3 | 0 | N |
| 12 | 79 | F | Control | Olfactory bulb | 4 | 0 | N |
|  |  |  |  | Olfactory tract (both) | 3 | 0 |  |
| 13 | 79 | F | Control | Olfactory bulb + tract (right) | 2 | 0 | N |
|  |  |  |  | Olfactory bulb + tract (left) | 1 | 0 |  |
| 14 | 79 | F | Control | Olfactory bulb | 1 | 0 | N |
|  |  |  |  | Olfactory bulb + tract | 4 | 0 |  |
| 15 | 80 | M | Control | Olfactory bulb + tract | 3 | 0 | N |
| 16 | 84 | F | Control | Olfactory bulb + tract (both) | 5 | 0 | N |
| 17 | 51 | M | COVID19+ | Olfactory bulb + tract (right) | 2 | 0 | N |
|  |  |  |  | Olfactory bulb + tract (left) | 5 | 0 |  |
| 18 | 52 | M | COVID19+ | Olfactory bulb | 3 | 0 | N |
|  |  |  |  | Olfactory tract | 4 | 0 |  |
| 19 | 56 | M | COVID19+ | Olfactory bulb + tract (right) | 1 | 0 | N |
|  |  |  |  | Olfactory bulb + tract (left) | 0 | 0 |  |
| 20 | 59 | M | COVID19+ | Olfactory bulb + tract (right) | 2 | 0 | N |
|  |  |  |  | Olfactory bulb + tract (left) | 5 | 0 |  |
| 21 | 61 | F | COVID19+ | Olfactory bulb + tract (right) | 2 | 0 | N |
|  |  |  |  | Olfactory bulb + tract (left) | 1 | 0 |  |
| 22 | 63 | F | COVID19+ | Olfactory bulb + tract (right) | 2 | 0 | N |
|  |  |  |  | Olfactory tract (left) | 1 | 0 |  |
| 23 | 73 | F | COVID19+ | Olfactory bulb + tract (right) | 1 | 0 | N |
|  |  |  |  | Olfactory bulb + tract (left) | 1 | 0 |  |
| 24 | 76 | M | COVID19+ | Olfactory bulb + tract (right) | 5 | 1 | N |
|  |  |  |  | Olfactory bulb + tract (left) | 2 | 0 |  |
| 25 | 76 | F | COVID19+ | Olfactory bulb + tract (right) | 3 | 0 | N |
|  |  |  |  | Olfactory bulb + tract (left) | 3 | 0 |  |
| 26 | 77 | F | COVID19+ | Olfactory tract (both) | 1 | 0 | N |
| 27 | 78 | M | COVID19+ | Olfactory bulb + tract (both) | 1 | 0 | N |
| 28 | 78 | F | COVID19+ | Olfactory bulb (right) | 2 | 0 | N |
|  |  |  |  | Olfactory bulb (left) | 2 | 0 |  |
| 29 | 79 | M | COVID19+ | Olfactory bulb + tract (right) | 4 | 0 | N |
|  |  |  |  | Olfactory bulb + tract (left) | 2 | 0 |  |
| 30 | 80 | F | COVID19+ | Olfactory bulb + tract (right) | 3 | 0 | N |
|  |  |  |  | Olfactory bulb + tract (left) | 2 | 0 |  |
| 31 | 83 | F | COVID19+ | Olfactory bulb + tract (right) | 2 | 2 | Y |
|  |  |  |  | Olfactory bulb + tract (left) | 3 | 2 |  |
| 32 | 84 | M | COVID19+ | Olfactory bulb + tract (both) | 4 | 2 | Y |
| 33 | 84 | M | COVID19+ | Olfactory bulb + tract (right) | 2 | 0 | N |
|  |  |  |  | Olfactory bulb + tract (left) | 2 | 0 |  |
| 34 | 85 | M | COVID19+ | Olfactory bulb + tract (right) | 2 | 3 | Y |
|  |  |  |  | Olfactory bulb + tract (left) | 3 | 3 |  |
| 35 | 86 | M | COVID19+ | Olfactory tract (right) | 2 | 0 | N |
|  |  |  |  | Olfactory bulb + tract (left) | 1 | 0 |  |
| 36 | 86 | F | COVID19+ | Olfactory bulb + tract (right) | 2 | 0 | N |
|  |  |  |  | Olfactory bulb + tract (left) | 2 | 0 |  |
| 37 | 89 | M | COVID19+ | Olfactory bulb + tract (right) | 3 | 0 | N |
|  |  |  |  | Olfactory bulb + tract (left) | 2 | 0 |  |
| 38 | 89 | M | COVID19+ | Olfactory bulb + tract (both) | 6 | 0 | N |
| 39 | 72 | M | LBD | Olfactory bulb + tract | 5 | 5 | Y |
| 40 | 73 | M | LBD | Olfactory tract | 0 | 0 | Y |
|  |  |  |  | Olfactory bulb | 3 | 4 |  |
| 41 | 77 | M | LBD | Olfactory bulb | 4 | 4 | N |
| 42 | 79 | M | LBD | Olfactory bulb + tract | 4 | 5 | Y |
| 43 | 82 | M | LBD | Olfactory bulb + tract | 2 | 5 | Y |
|  |  |  |  | Olfactory bulb | 2 | 5 |  |
| 44 | 90 | M | LBD | Olfactory tract | 3 | 3 | Y |
| 45 | 52 | F | AD | Olfactory bulb + tract (both) | 7 | 5 | Y |
| 46 | 78 | F | AD | Olfactory bulb + tract | 3 | 0 | N |
|  |  |  |  | Olfactory bulb + tract | 3 | 0 |  |
| 47 | 79 | M | AD | Olfactory bulb + tract | 2 | 0 | N |
|  |  |  |  | Olfactory bulb + tract | 3 | 0 |  |
| 48 | 74 | M | OND | Olfactory bulb + tract | 1 | 0 | N |
| 49 | 79 | M | OND | Olfactory tract (both) | 2 | 2 | Y |
| 50 | 81 | M | OND | Olfactory tract | 1 | 0 | N |

AON, anterior olfactory nucleus; SN, substantia nigra; DNV, dorsal motor nucleus of the vagus; LBD, Lewy Body disease; AD, Alzheimer's disease; OND, other neurodegenerative disease

**Supplemental Table 4:**
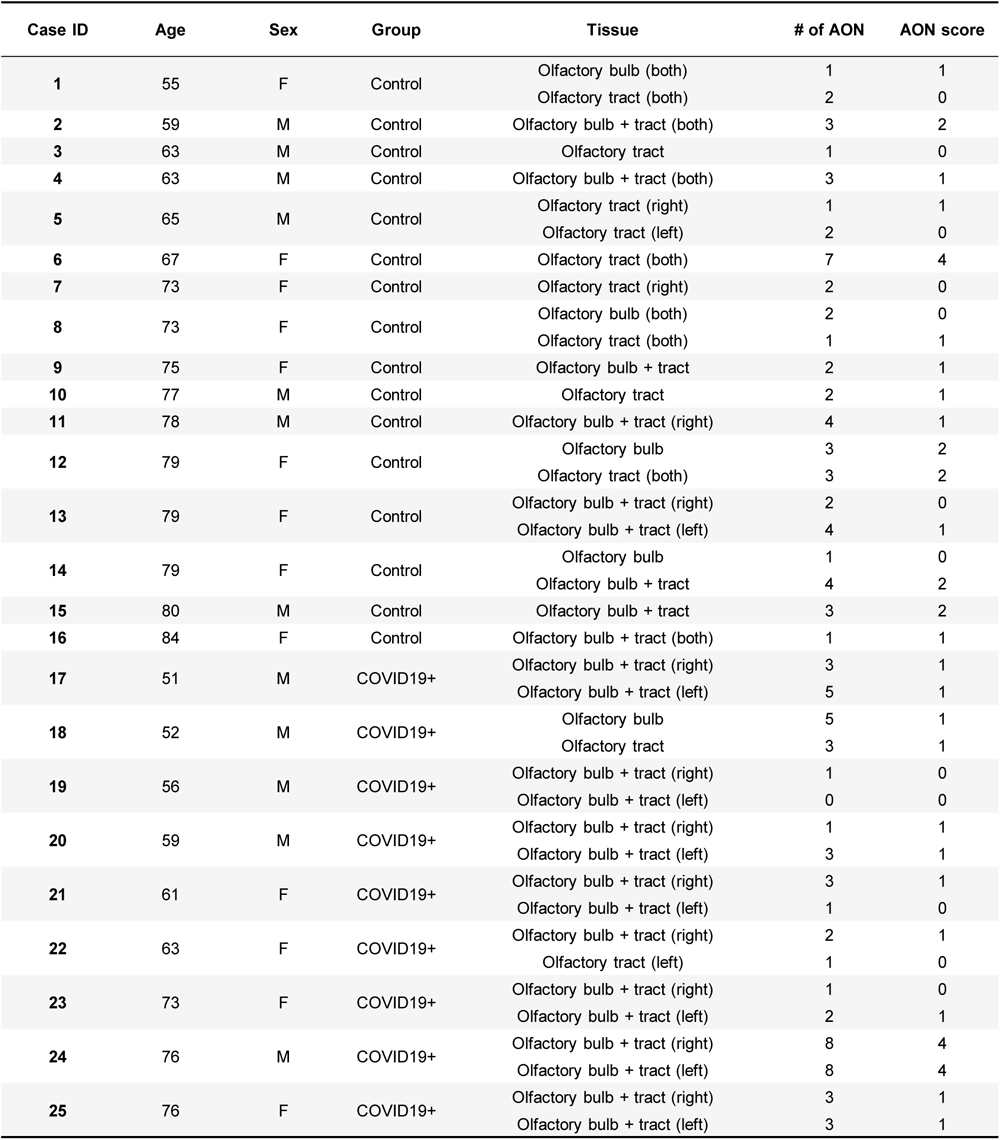
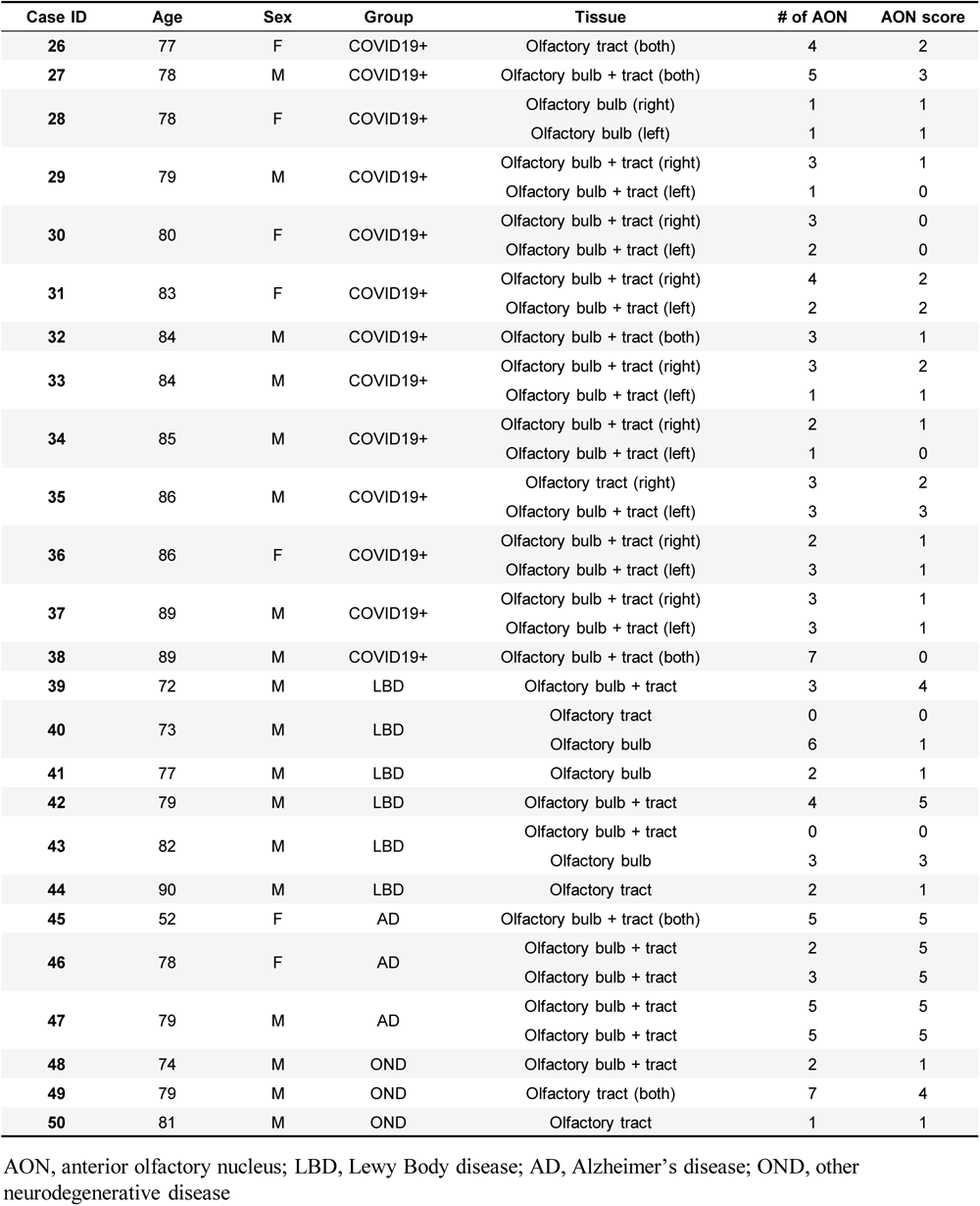
Phosphorylated tau staining.

| Case ID | Age | Sex | Group | Tissue | # of AON | AON score |
| --- | --- | --- | --- | --- | --- | --- |
| 1 | 55 | F | Control | Olfactory bulb (both) | 1 | 1 |
|  |  |  |  | Olfactory tract (both) | 2 | 0 |
| 2 | 59 | M | Control | Olfactory bulb + tract (both) | 3 | 2 |
| 3 | 63 | M | Control | Olfactory tract | 1 | 0 |
| 4 | 63 | M | Control | Olfactory bulb + tract (both) | 3 | 1 |
| 5 | 65 | M | Control | Olfactory tract (right) | 1 | 1 |
|  |  |  |  | Olfactory tract (left) | 2 | 0 |
| 6 | 67 | F | Control | Olfactory tract (both) | 7 | 4 |
| 7 | 73 | F | Control | Olfactory tract (right) | 2 | 0 |
| 8 | 73 | F | Control | Olfactory bulb (both) | 2 | 0 |
|  |  |  |  | Olfactory tract (both) | 1 | 1 |
| 9 | 75 | F | Control | Olfactory bulb + tract | 2 | 1 |
| 10 | 77 | M | Control | Olfactory tract | 2 | 1 |
| 11 | 78 | M | Control | Olfactory bulb + tract (right) | 4 | 1 |
| 12 | 79 | F | Control | Olfactory bulb | 3 | 2 |
|  |  |  |  | Olfactory tract (both) | 3 | 2 |
| 13 | 79 | F | Control | Olfactory bulb + tract (right) | 2 | 0 |
|  |  |  |  | Olfactory bulb + tract (left) | 4 | 1 |
| 14 | 79 | F | Control | Olfactory bulb | 1 | 0 |
|  |  |  |  | Olfactory bulb + tract | 4 | 2 |
| 15 | 80 | M | Control | Olfactory bulb + tract | 3 | 2 |
| 16 | 84 | F | Control | Olfactory bulb + tract (both) | 1 | 1 |
| 17 | 51 | M | COVID19+ | Olfactory bulb + tract (right) | 3 | 1 |
|  |  |  |  | Olfactory bulb + tract (left) | 5 | 1 |
| 18 | 52 | M | COVID19+ | Olfactory bulb | 5 | 1 |
|  |  |  |  | Olfactory tract | 3 | 1 |
| 19 | 56 | M | COVID19+ | Olfactory bulb + tract (right) | 1 | 0 |
|  |  |  |  | Olfactory bulb + tract (left) | 0 | 0 |
| 20 | 59 | M | COVID19+ | Olfactory bulb + tract (right) | 1 | 1 |
|  |  |  |  | Olfactory bulb + tract (left) | 3 | 1 |
| 21 | 61 | F | COVID19+ | Olfactory bulb + tract (right) | 3 | 1 |
|  |  |  |  | Olfactory bulb + tract (left) | 1 | 0 |
| 22 | 63 | F | COVID19+ | Olfactory bulb + tract (right) | 2 | 1 |
|  |  |  |  | Olfactory tract (left) | 1 | 0 |
| 23 | 73 | F | COVID19+ | Olfactory bulb + tract (right) | 1 | 0 |
|  |  |  |  | Olfactory bulb + tract (left) | 2 | 1 |
| 24 | 76 | M | COVID19+ | Olfactory bulb + tract (right) | 8 | 4 |
|  |  |  |  | Olfactory bulb + tract (left) | 8 | 4 |
| 25 | 76 | F | COVID19+ | Olfactory bulb + tract (right) | 3 | 1 |
|  |  |  |  | Olfactory bulb + tract (left) | 3 | 1 |
| 26 | 77 | F | COVID19+ | Olfactory tract (both) | 4 | 2 |
| 27 | 78 | M | COVID19+ | Olfactory bulb + tract (both) | 5 | 3 |
| 28 | 78 | F | COVID19+ | Olfactory bulb (right) | 1 | 1 |
|  |  |  |  | Olfactory bulb (left) | 1 | 1 |
| 29 | 79 | M | COVID19+ | Olfactory bulb + tract (right) | 3 | 1 |
|  |  |  |  | Olfactory bulb + tract (left) | 1 | 0 |
| 30 | 80 | F | COVID19+ | Olfactory bulb + tract (right) | 3 | 0 |
|  |  |  |  | Olfactory bulb + tract (left) | 2 | 0 |
| 31 | 83 | F | COVID19+ | Olfactory bulb + tract (right) | 4 | 2 |
|  |  |  |  | Olfactory bulb + tract (left) | 2 | 2 |
| 32 | 84 | M | COVID19+ | Olfactory bulb + tract (both) | 3 | 1 |
| 33 | 84 | M | COVID19+ | Olfactory bulb + tract (right) | 3 | 2 |
|  |  |  |  | Olfactory bulb + tract (left) | 1 | 1 |
| 34 | 85 | M | COVID19+ | Olfactory bulb + tract (right) | 2 | 1 |
|  |  |  |  | Olfactory bulb + tract (left) | 1 | 0 |
| 35 | 86 | M | COVID19+ | Olfactory tract (right) | 3 | 2 |
|  |  |  |  | Olfactory bulb + tract (left) | 3 | 3 |
| 36 | 86 | F | COVID19+ | Olfactory bulb + tract (right) | 2 | 1 |
|  |  |  |  | Olfactory bulb + tract (left) | 3 | 1 |
| 37 | 89 | M | COVID19+ | Olfactory bulb + tract (right) | 3 | 1 |
|  |  |  |  | Olfactory bulb + tract (left) | 3 | 1 |
| 38 | 89 | M | COVID19+ | Olfactory bulb + tract (both) | 7 | 0 |
| 39 | 72 | M | LBD | Olfactory bulb + tract | 3 | 4 |
| 40 | 73 | M | LBD | Olfactory tract | 0 | 0 |
|  |  |  |  | Olfactory bulb | 6 | 1 |
| 41 | 77 | M | LBD | Olfactory bulb | 2 | 1 |
| 42 | 79 | M | LBD | Olfactory bulb + tract | 4 | 5 |
| 43 | 82 | M | LBD | Olfactory bulb + tract | 0 | 0 |
|  |  |  |  | Olfactory bulb | 3 | 3 |
| 44 | 90 | M | LBD | Olfactory tract | 2 | 1 |
| 45 | 52 | F | AD | Olfactory bulb + tract (both) | 5 | 5 |
| 46 | 78 | F | AD | Olfactory bulb + tract | 2 | 5 |
|  |  |  |  | Olfactory bulb + tract | 3 | 5 |
| 47 | 79 | M | AD | Olfactory bulb + tract | 5 | 5 |
|  |  |  |  | Olfactory bulb + tract | 5 | 5 |
| 48 | 74 | M | OND | Olfactory bulb + tract | 2 | 1 |
| 49 | 79 | M | OND | Olfactory tract (both) | 7 | 4 |
| 50 | 81 | M | OND | Olfactory tract | 1 | 1 |

AON, anterior olfactory nucleus; LBD, Lewy Body disease; AD, Alzheimer's disease; OND, other neurodegenerative disease

**Supplemental Table 5:**
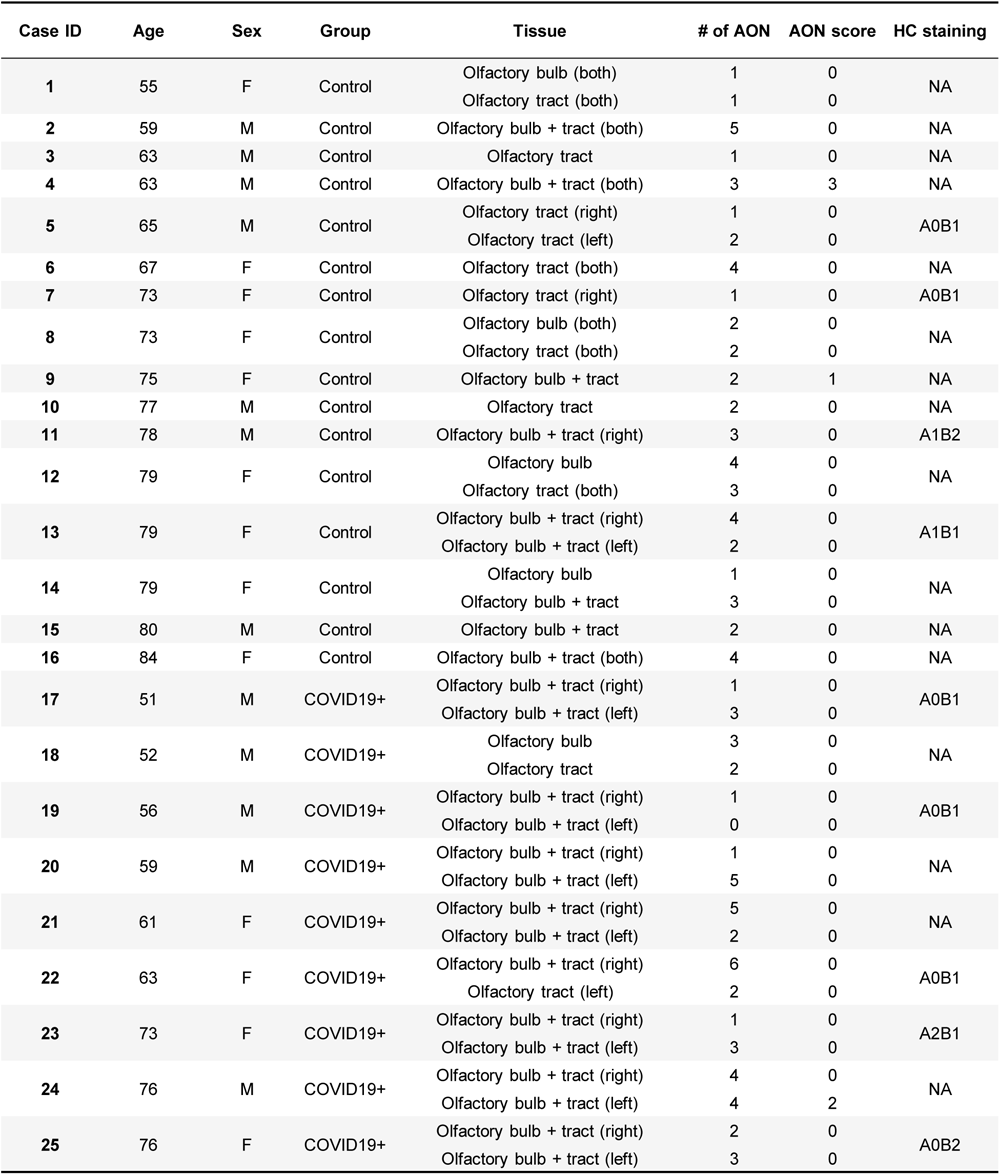
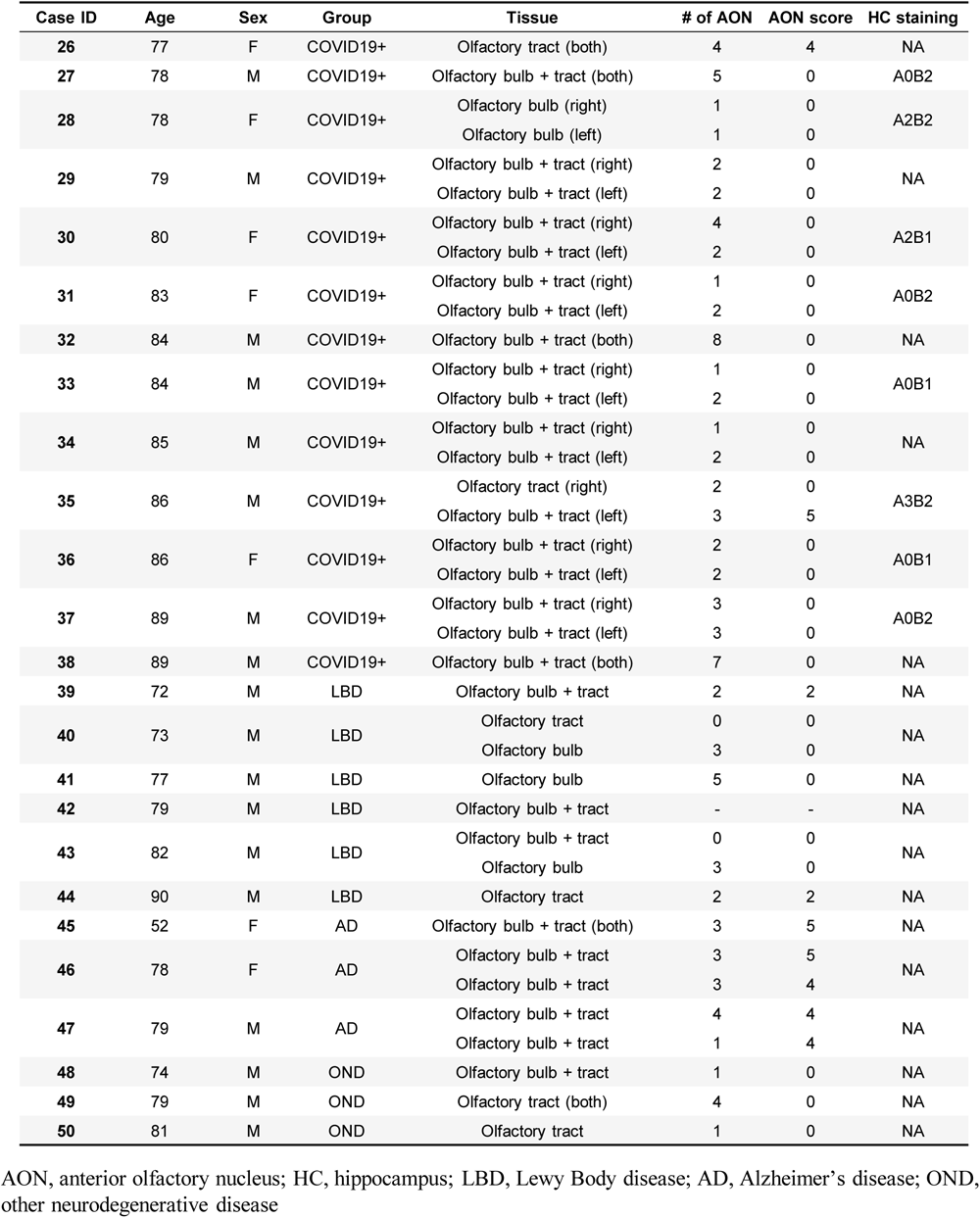
Amyloid-β staining.

| Case ID | Age | Sex | Group | Tissue | # of AON | AON score | HC staining |
| --- | --- | --- | --- | --- | --- | --- | --- |
| 1 | 55 | F | Control | Olfactory bulb (both) | 1 | 0 | NA |
|  |  |  |  | Olfactory tract (both) | 1 | 0 |  |
| 2 | 59 | M | Control | Olfactory bulb + tract (both) | 5 | 0 | NA |
| 3 | 63 | M | Control | Olfactory tract | 1 | 0 | NA |
| 4 | 63 | M | Control | Olfactory bulb + tract (both) | 3 | 3 | NA |
| 5 | 65 | M | Control | Olfactory tract (right) | 1 | 0 | A0B1 |
|  |  |  |  | Olfactory tract (left) | 2 | 0 |  |
| 6 | 67 | F | Control | Olfactory tract (both) | 4 | 0 | NA |
| 7 | 73 | F | Control | Olfactory tract (right) | 1 | 0 | A0B1 |
| 8 | 73 | F | Control | Olfactory bulb (both) | 2 | 0 | NA |
|  |  |  |  | Olfactory tract (both) | 2 | 0 |  |
| 9 | 75 | F | Control | Olfactory bulb + tract | 2 | 1 | NA |
| 10 | 77 | M | Control | Olfactory tract | 2 | 0 | NA |
| 11 | 78 | M | Control | Olfactory bulb + tract (right) | 3 | 0 | A1B2 |
| 12 | 79 | F | Control | Olfactory bulb | 4 | 0 | NA |
|  |  |  |  | Olfactory tract (both) | 3 | 0 |  |
| 13 | 79 | F | Control | Olfactory bulb + tract (right) | 4 | 0 | A1B1 |
|  |  |  |  | Olfactory bulb + tract (left) | 2 | 0 |  |
| 14 | 79 | F | Control | Olfactory bulb | 1 | 0 | NA |
|  |  |  |  | Olfactory bulb + tract | 3 | 0 |  |
| 15 | 80 | M | Control | Olfactory bulb + tract | 2 | 0 | NA |
| 16 | 84 | F | Control | Olfactory bulb + tract (both) | 4 | 0 | NA |
| 17 | 51 | M | COVID19+ | Olfactory bulb + tract (right) | 1 | 0 | A0B1 |
|  |  |  |  | Olfactory bulb + tract (left) | 3 | 0 |  |
| 18 | 52 | M | COVID19+ | Olfactory bulb | 3 | 0 | NA |
|  |  |  |  | Olfactory tract | 2 | 0 |  |
| 19 | 56 | M | COVID19+ | Olfactory bulb + tract (right) | 1 | 0 | A0B1 |
|  |  |  |  | Olfactory bulb + tract (left) | 0 | 0 |  |
| 20 | 59 | M | COVID19+ | Olfactory bulb + tract (right) | 1 | 0 | NA |
|  |  |  |  | Olfactory bulb + tract (left) | 5 | 0 |  |
| 21 | 61 | F | COVID19+ | Olfactory bulb + tract (right) | 5 | 0 | NA |
|  |  |  |  | Olfactory bulb + tract (left) | 2 | 0 |  |
| 22 | 63 | F | COVID19+ | Olfactory bulb + tract (right) | 6 | 0 | A0B1 |
|  |  |  |  | Olfactory tract (left) | 2 | 0 |  |
| 23 | 73 | F | COVID19+ | Olfactory bulb + tract (right) | 1 | 0 | A2B1 |
|  |  |  |  | Olfactory bulb + tract (left) | 3 | 0 |  |
| 24 | 76 | M | COVID19+ | Olfactory bulb + tract (right) | 4 | 0 | NA |
|  |  |  |  | Olfactory bulb + tract (left) | 4 | 2 |  |
| 25 | 76 | F | COVID19+ | Olfactory bulb + tract (right) | 2 | 0 | A0B2 |
|  |  |  |  | Olfactory bulb + tract (left) | 3 | 0 |  |
| 26 | 77 | F | COVID19+ | Olfactory tract (both) | 4 | 4 | NA |
| 27 | 78 | M | COVID19+ | Olfactory bulb + tract (both) | 5 | 0 | A0B2 |
| 28 | 78 | F | COVID19+ | Olfactory bulb (right) | 1 | 0 | A2B2 |
|  |  |  |  | Olfactory bulb (left) | 1 | 0 |  |
| 29 | 79 | M | COVID19+ | Olfactory bulb + tract (right) | 2 | 0 | NA |
|  |  |  |  | Olfactory bulb + tract (left) | 2 | 0 |  |
| 30 | 80 | F | COVID19+ | Olfactory bulb + tract (right) | 4 | 0 | A2B1 |
|  |  |  |  | Olfactory bulb + tract (left) | 2 | 0 |  |
| 31 | 83 | F | COVID19+ | Olfactory bulb + tract (right) | 1 | 0 | A0B2 |
|  |  |  |  | Olfactory bulb + tract (left) | 2 | 0 |  |
| 32 | 84 | M | COVID19+ | Olfactory bulb + tract (both) | 8 | 0 | NA |
| 33 | 84 | M | COVID19+ | Olfactory bulb + tract (right) | 1 | 0 | A0B1 |
|  |  |  |  | Olfactory bulb + tract (left) | 2 | 0 |  |
| 34 | 85 | M | COVID19+ | Olfactory bulb + tract (right) | 1 | 0 | NA |
|  |  |  |  | Olfactory bulb + tract (left) | 2 | 0 |  |
| 35 | 86 | M | COVID19+ | Olfactory tract (right) | 2 | 0 | A3B2 |
|  |  |  |  | Olfactory bulb + tract (left) | 3 | 5 |  |
| 36 | 86 | F | COVID19+ | Olfactory bulb + tract (right) | 2 | 0 | A0B1 |
|  |  |  |  | Olfactory bulb + tract (left) | 2 | 0 |  |
| 37 | 89 | M | COVID19+ | Olfactory bulb + tract (right) | 3 | 0 | A0B2 |
|  |  |  |  | Olfactory bulb + tract (left) | 3 | 0 |  |
| 38 | 89 | M | COVID19+ | Olfactory bulb + tract (both) | 7 | 0 | NA |
| 39 | 72 | M | LBD | Olfactory bulb + tract | 2 | 2 | NA |
| 40 | 73 | M | LBD | Olfactory tract | 0 | 0 | NA |
|  |  |  |  | Olfactory bulb | 3 | 0 |  |
| 41 | 77 | M | LBD | Olfactory bulb | 5 | 0 | NA |
| 42 | 79 | M | LBD | Olfactory bulb + tract | - | - | NA |
| 43 | 82 | M | LBD | Olfactory bulb + tract | 0 | 0 | NA |
|  |  |  |  | Olfactory bulb | 3 | 0 |  |
| 44 | 90 | M | LBD | Olfactory tract | 2 | 2 | NA |
| 45 | 52 | F | AD | Olfactory bulb + tract (both) | 3 | 5 | NA |
| 46 | 78 | F | AD | Olfactory bulb + tract | 3 | 5 | NA |
|  |  |  |  | Olfactory bulb + tract | 3 | 4 |  |
| 47 | 79 | M | AD | Olfactory bulb + tract | 4 | 4 | NA |
|  |  |  |  | Olfactory bulb + tract | 1 | 4 |  |
| 48 | 74 | M | OND | Olfactory bulb + tract | 1 | 0 | NA |
| 49 | 79 | M | OND | Olfactory tract (both) | 4 | 0 | NA |
| 50 | 81 | M | OND | Olfactory tract | 1 | 0 | NA |

AON, anterior olfactory nucleus; HC, hippocampus; LBD, Lewy Body disease; AD, Alzheimer's disease; OND, other neurodegenerative disease

**Supplemental Table 6:**
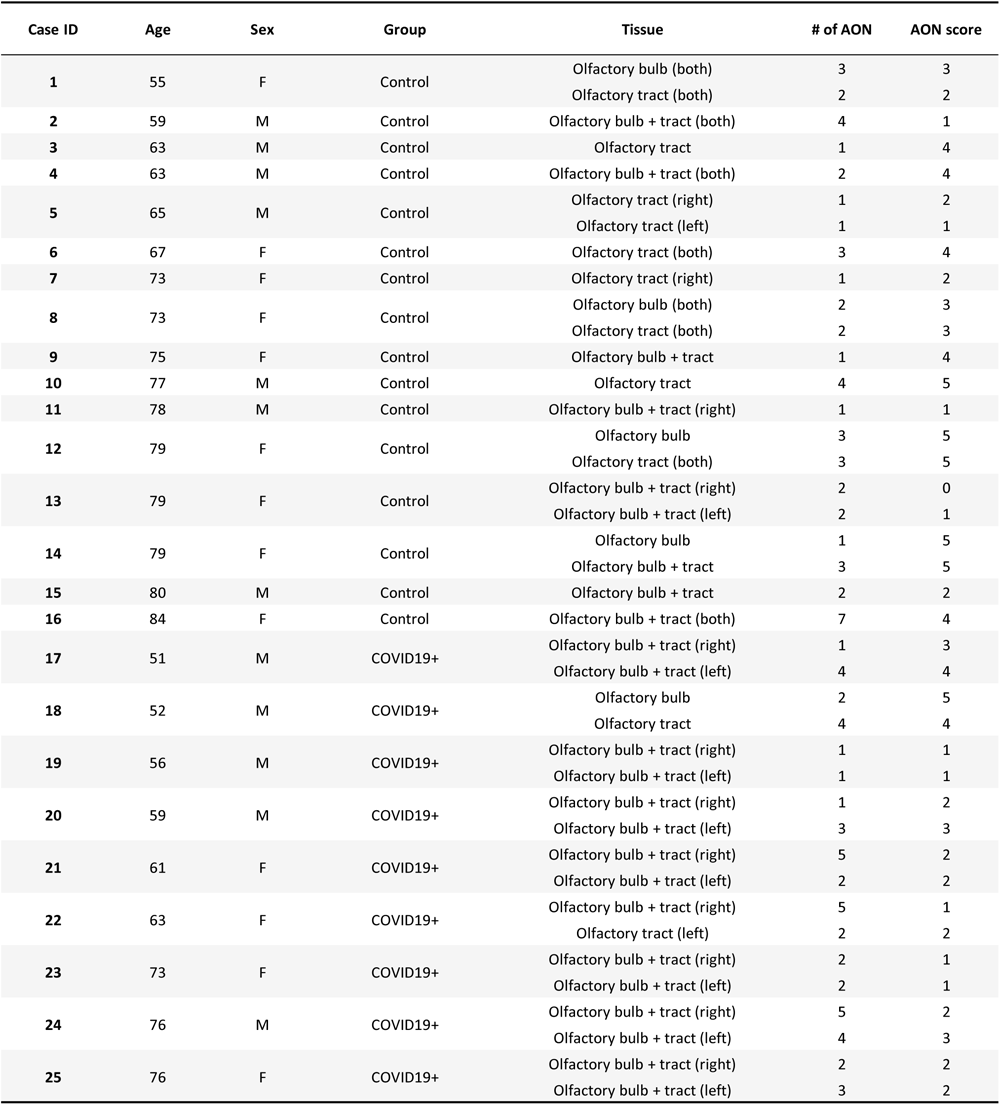
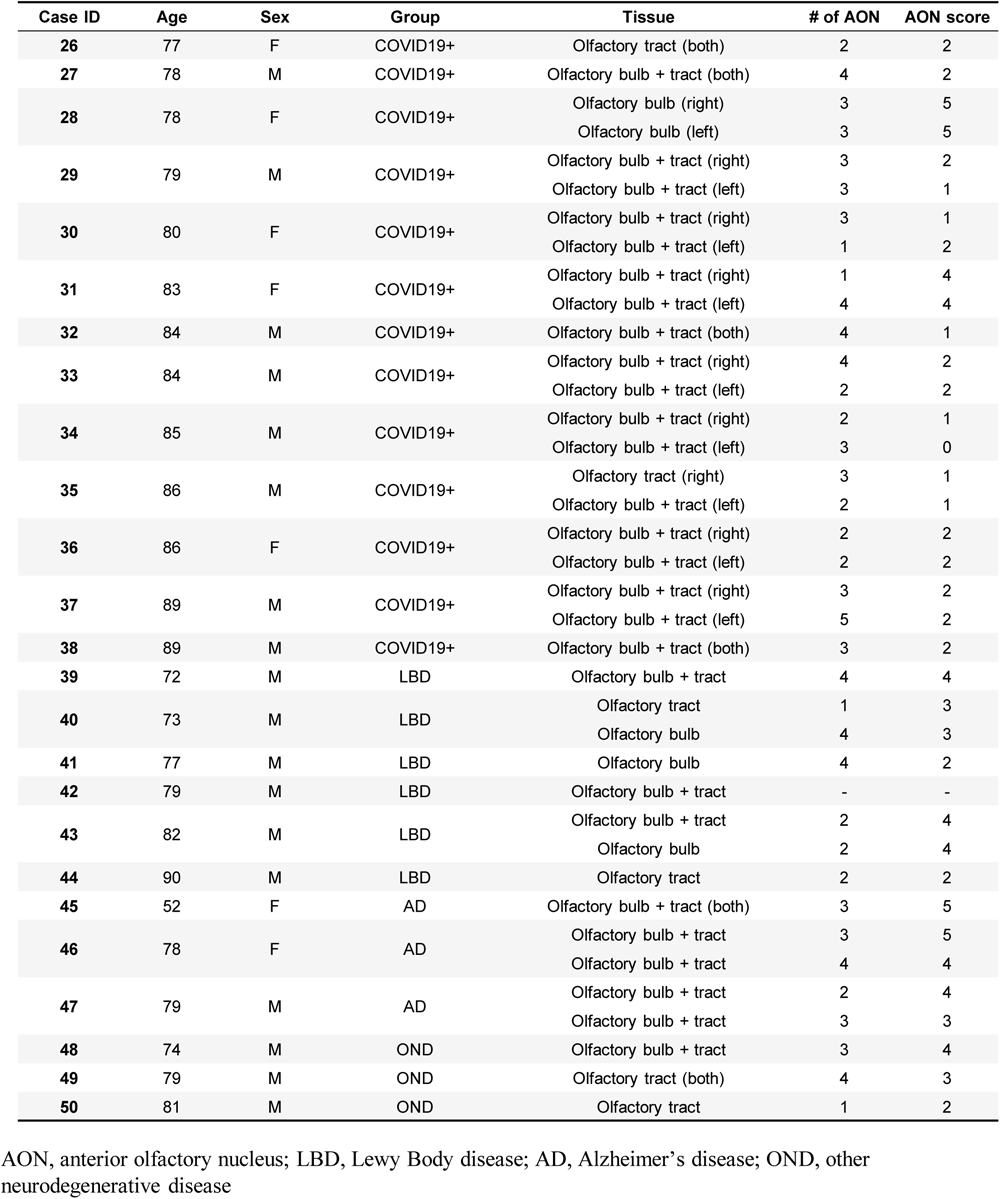
CD68 staining.

| Case ID | Age | Sex | Group | Tissue | # of AON | AON score |
| --- | --- | --- | --- | --- | --- | --- |
| 1 | 55 | F | Control | Olfactory bulb (both) | 3 | 3 |
|  |  |  |  | Olfactory tract (both) | 2 | 2 |
| 2 | 59 | M | Control | Olfactory bulb + tract (both) | 4 | 1 |
| 3 | 63 | M | Control | Olfactory tract | 1 | 4 |
| 4 | 63 | M | Control | Olfactory bulb + tract (both) | 2 | 4 |
| 5 | 65 | M | Control | Olfactory tract (right) | 1 | 2 |
|  |  |  |  | Olfactory tract (left) | 1 | 1 |
| 6 | 67 | F | Control | Olfactory tract (both) | 3 | 4 |
| 7 | 73 | F | Control | Olfactory tract (right) | 1 | 2 |
| 8 | 73 | F | Control | Olfactory bulb (both) | 2 | 3 |
|  |  |  |  | Olfactory tract (both) | 2 | 3 |
| 9 | 75 | F | Control | Olfactory bulb + tract | 1 | 4 |
| 10 | 77 | M | Control | Olfactory tract | 4 | 5 |
| 11 | 78 | M | Control | Olfactory bulb + tract (right) | 1 | 1 |
| 12 | 79 | F | Control | Olfactory bulb | 3 | 5 |
|  |  |  |  | Olfactory tract (both) | 3 | 5 |
| 13 | 79 | F | Control | Olfactory bulb + tract (right) | 2 | 0 |
|  |  |  |  | Olfactory bulb + tract (left) | 2 | 1 |
| 14 | 79 | F | Control | Olfactory bulb | 1 | 5 |
|  |  |  |  | Olfactory bulb + tract | 3 | 5 |
| 15 | 80 | M | Control | Olfactory bulb + tract | 2 | 2 |
| 16 | 84 | F | Control | Olfactory bulb + tract (both) | 7 | 4 |
| 17 | 51 | M | COVID19+ | Olfactory bulb + tract (right) | 1 | 3 |
|  |  |  |  | Olfactory bulb + tract (left) | 4 | 4 |
| 18 | 52 | M | COVID19+ | Olfactory bulb | 2 | 5 |
|  |  |  |  | Olfactory tract | 4 | 4 |
| 19 | 56 | M | COVID19+ | Olfactory bulb + tract (right) | 1 | 1 |
|  |  |  |  | Olfactory bulb + tract (left) | 1 | 1 |
| 20 | 59 | M | COVID19+ | Olfactory bulb + tract (right) | 1 | 2 |
|  |  |  |  | Olfactory bulb + tract (left) | 3 | 3 |
| 21 | 61 | F | COVID19+ | Olfactory bulb + tract (right) | 5 | 2 |
|  |  |  |  | Olfactory bulb + tract (left) | 2 | 2 |
| 22 | 63 | F | COVID19+ | Olfactory bulb + tract (right) | 5 | 1 |
|  |  |  |  | Olfactory tract (left) | 2 | 2 |
| 23 | 73 | F | COVID19+ | Olfactory bulb + tract (right) | 2 | 1 |
|  |  |  |  | Olfactory bulb + tract (left) | 2 | 1 |
| 24 | 76 | M | COVID19+ | Olfactory bulb + tract (right) | 5 | 2 |
|  |  |  |  | Olfactory bulb + tract (left) | 4 | 3 |
| 25 | 76 | F | COVID19+ | Olfactory bulb + tract (right) | 2 | 2 |
|  |  |  |  | Olfactory bulb + tract (left) | 3 | 2 |
| 26 | 77 | F | COVID19+ | Olfactory tract (both) | 2 | 2 |
| 27 | 78 | M | COVID19+ | Olfactory bulb + tract (both) | 4 | 2 |
| 28 | 78 | F | COVID19+ | Olfactory bulb (right) | 3 | 5 |
|  |  |  |  | Olfactory bulb (left) | 3 | 5 |
| 29 | 79 | M | COVID19+ | Olfactory bulb + tract (right) | 3 | 2 |
|  |  |  |  | Olfactory bulb + tract (left) | 3 | 1 |
| 30 | 80 | F | COVID19+ | Olfactory bulb + tract (right) | 3 | 1 |
|  |  |  |  | Olfactory bulb + tract (left) | 1 | 2 |
| 31 | 83 | F | COVID19+ | Olfactory bulb + tract (right) | 1 | 4 |
|  |  |  |  | Olfactory bulb + tract (left) | 4 | 4 |
| 32 | 84 | M | COVID19+ | Olfactory bulb + tract (both) | 4 | 1 |
| 33 | 84 | M | COVID19+ | Olfactory bulb + tract (right) | 4 | 2 |
|  |  |  |  | Olfactory bulb + tract (left) | 2 | 2 |
| 34 | 85 | M | COVID19+ | Olfactory bulb + tract (right) | 2 | 1 |
|  |  |  |  | Olfactory bulb + tract (left) | 3 | 0 |
| 35 | 86 | M | COVID19+ | Olfactory tract (right) | 3 | 1 |
|  |  |  |  | Olfactory bulb + tract (left) | 2 | 1 |
| 36 | 86 | F | COVID19+ | Olfactory bulb + tract (right) | 2 | 2 |
|  |  |  |  | Olfactory bulb + tract (left) | 2 | 2 |
| 37 | 89 | M | COVID19+ | Olfactory bulb + tract (right) | 3 | 2 |
|  |  |  |  | Olfactory bulb + tract (left) | 5 | 2 |
| 38 | 89 | M | COVID19+ | Olfactory bulb + tract (both) | 3 | 2 |
| 39 | 72 | M | LBD | Olfactory bulb + tract | 4 | 4 |
| 40 | 73 | M | LBD | Olfactory tract | 1 | 3 |
|  |  |  |  | Olfactory bulb | 4 | 3 |
| 41 | 77 | M | LBD | Olfactory bulb | 4 | 2 |
| 42 | 79 | M | LBD | Olfactory bulb + tract | - | - |
| 43 | 82 | M | LBD | Olfactory bulb + tract | 2 | 4 |
|  |  |  |  | Olfactory bulb | 2 | 4 |
| 44 | 90 | M | LBD | Olfactory tract | 2 | 2 |
| 45 | 52 | F | AD | Olfactory bulb + tract (both) | 3 | 5 |
| 46 | 78 | F | AD | Olfactory bulb + tract | 3 | 5 |
|  |  |  |  | Olfactory bulb + tract | 4 | 4 |
| 47 | 79 | M | AD | Olfactory bulb + tract | 2 | 4 |
|  |  |  |  | Olfactory bulb + tract | 3 | 3 |
| 48 | 74 | M | OND | Olfactory bulb + tract | 3 | 4 |
| 49 | 79 | M | OND | Olfactory tract (both) | 4 | 3 |
| 50 | 81 | M | OND | Olfactory tract | 1 | 2 |

AON, anterior olfactory nucleus; LBD, Lewy Body disease; AD, Alzheimer's disease; OND, other neurodegenerative disease

**Supplemental Table 7:**
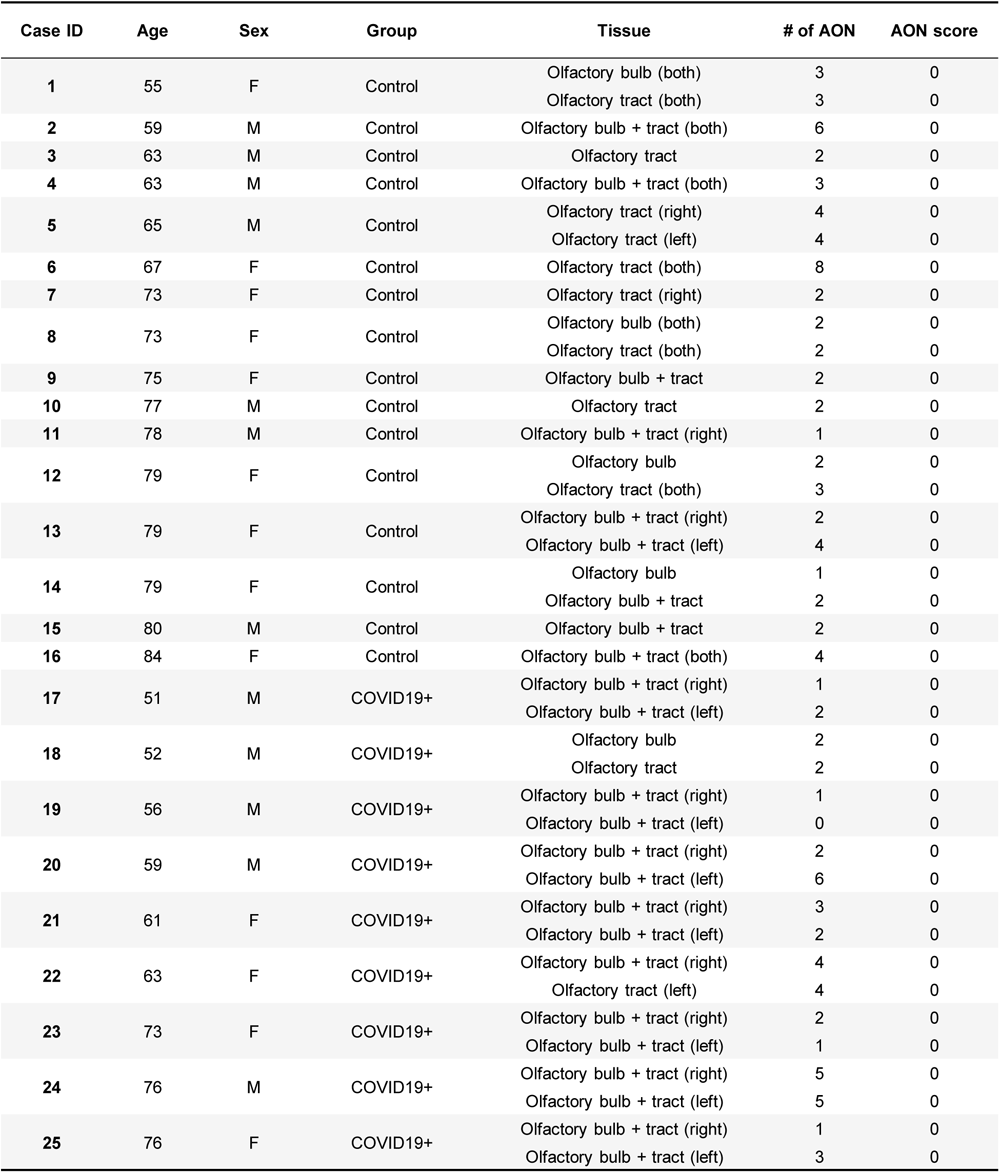
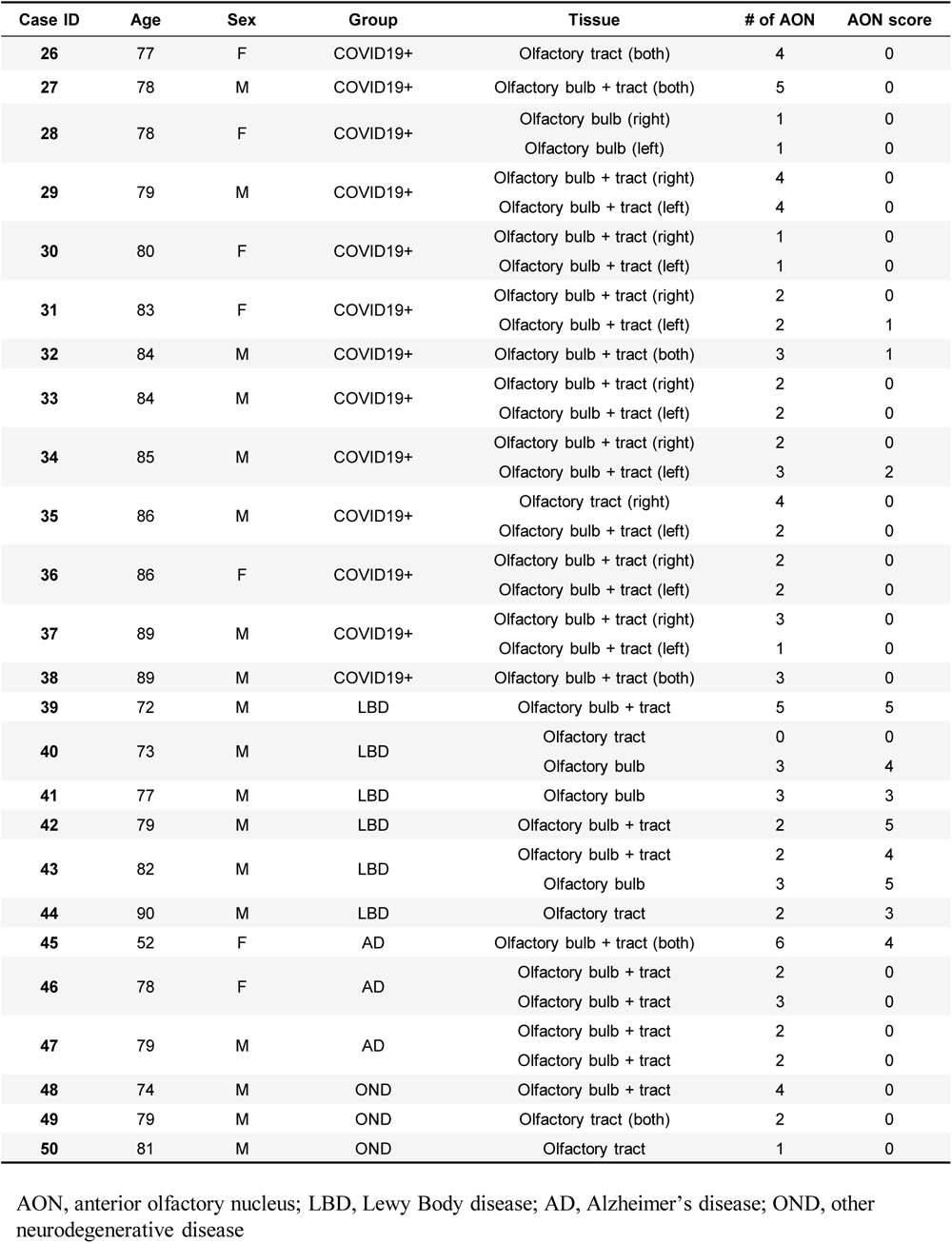
LB509 staining.

| Case ID | Age | Sex | Group | Tissue | # of AON | AON score |
| --- | --- | --- | --- | --- | --- | --- |
| 1 | 55 | F | Control | Olfactory bulb (both) | 3 | 0 |
|  |  |  |  | Olfactory tract (both) | 3 | 0 |
| 2 | 59 | M | Control | Olfactory bulb + tract (both) | 6 | 0 |
| 3 | 63 | M | Control | Olfactory tract | 2 | 0 |
| 4 | 63 | M | Control | Olfactory bulb + tract (both) | 3 | 0 |
| 5 | 65 | M | Control | Olfactory tract (right) | 4 | 0 |
|  |  |  |  | Olfactory tract (left) | 4 | 0 |
| 6 | 67 | F | Control | Olfactory tract (both) | 8 | 0 |
| 7 | 73 | F | Control | Olfactory tract (right) | 2 | 0 |
| 8 | 73 | F | Control | Olfactory bulb (both) | 2 | 0 |
|  |  |  |  | Olfactory tract (both) | 2 | 0 |
| 9 | 75 | F | Control | Olfactory bulb + tract | 2 | 0 |
| 10 | 77 | M | Control | Olfactory tract | 2 | 0 |
| 11 | 78 | M | Control | Olfactory bulb + tract (right) | 1 | 0 |
| 12 | 79 | F | Control | Olfactory bulb | 2 | 0 |
|  |  |  |  | Olfactory tract (both) | 3 | 0 |
| 13 | 79 | F | Control | Olfactory bulb + tract (right) | 2 | 0 |
|  |  |  |  | Olfactory bulb + tract (left) | 4 | 0 |
| 14 | 79 | F | Control | Olfactory bulb | 1 | 0 |
|  |  |  |  | Olfactory bulb + tract | 2 | 0 |
| 15 | 80 | M | Control | Olfactory bulb + tract | 2 | 0 |
| 16 | 84 | F | Control | Olfactory bulb + tract (both) | 4 | 0 |
| 17 | 51 | M | COVID19+ | Olfactory bulb + tract (right) | 1 | 0 |
|  |  |  |  | Olfactory bulb + tract (left) | 2 | 0 |
| 18 | 52 | M | COVID19+ | Olfactory bulb | 2 | 0 |
|  |  |  |  | Olfactory tract | 2 | 0 |
| 19 | 56 | M | COVID19+ | Olfactory bulb + tract (right) | 1 | 0 |
|  |  |  |  | Olfactory bulb + tract (left) | 0 | 0 |
| 20 | 59 | M | COVID19+ | Olfactory bulb + tract (right) | 2 | 0 |
|  |  |  |  | Olfactory bulb + tract (left) | 6 | 0 |
| 21 | 61 | F | COVID19+ | Olfactory bulb + tract (right) | 3 | 0 |
|  |  |  |  | Olfactory bulb + tract (left) | 2 | 0 |
| 22 | 63 | F | COVID19+ | Olfactory bulb + tract (right) | 4 | 0 |
|  |  |  |  | Olfactory tract (left) | 4 | 0 |
| 23 | 73 | F | COVID19+ | Olfactory bulb + tract (right) | 2 | 0 |
|  |  |  |  | Olfactory bulb + tract (left) | 1 | 0 |
| 24 | 76 | M | COVID19+ | Olfactory bulb + tract (right) | 5 | 0 |
|  |  |  |  | Olfactory bulb + tract (left) | 5 | 0 |
| 25 | 76 | F | COVID19+ | Olfactory bulb + tract (right) | 1 | 0 |
|  |  |  |  | Olfactory bulb + tract (left) | 3 | 0 |
| 26 | 77 | F | COVID19+ | Olfactory tract (both) | 4 | 0 |
| 27 | 78 | M | COVID19+ | Olfactory bulb + tract (both) | 5 | 0 |
| 28 | 78 | F | COVID19+ | Olfactory bulb (right) | 1 | 0 |
|  |  |  |  | Olfactory bulb (left) | 1 | 0 |
| 29 | 79 | M | COVID19+ | Olfactory bulb + tract (right) | 4 | 0 |
|  |  |  |  | Olfactory bulb + tract (left) | 4 | 0 |
| 30 | 80 | F | COVID19+ | Olfactory bulb + tract (right) | 1 | 0 |
|  |  |  |  | Olfactory bulb + tract (left) | 1 | 0 |
| 31 | 83 | F | COVID19+ | Olfactory bulb + tract (right) | 2 | 0 |
|  |  |  |  | Olfactory bulb + tract (left) | 2 | 1 |
| 32 | 84 | M | COVID19+ | Olfactory bulb + tract (both) | 3 | 1 |
| 33 | 84 | M | COVID19+ | Olfactory bulb + tract (right) | 2 | 0 |
|  |  |  |  | Olfactory bulb + tract (left) | 2 | 0 |
| 34 | 85 | M | COVID19+ | Olfactory bulb + tract (right) | 2 | 0 |
|  |  |  |  | Olfactory bulb + tract (left) | 3 | 2 |
| 35 | 86 | M | COVID19+ | Olfactory tract (right) | 4 | 0 |
|  |  |  |  | Olfactory bulb + tract (left) | 2 | 0 |
| 36 | 86 | F | COVID19+ | Olfactory bulb + tract (right) | 2 | 0 |
|  |  |  |  | Olfactory bulb + tract (left) | 2 | 0 |
| 37 | 89 | M | COVID19+ | Olfactory bulb + tract (right) | 3 | 0 |
|  |  |  |  | Olfactory bulb + tract (left) | 1 | 0 |
| 38 | 89 | M | COVID19+ | Olfactory bulb + tract (both) | 3 | 0 |
| 39 | 72 | M | LBD | Olfactory bulb + tract | 5 | 5 |
| 40 | 73 | M | LBD | Olfactory tract | 0 | 0 |
|  |  |  |  | Olfactory bulb | 3 | 4 |
| 41 | 77 | M | LBD | Olfactory bulb | 3 | 3 |
| 42 | 79 | M | LBD | Olfactory bulb + tract | 2 | 5 |
| 43 | 82 | M | LBD | Olfactory bulb + tract | 2 | 4 |
|  |  |  |  | Olfactory bulb | 3 | 5 |
| 44 | 90 | M | LBD | Olfactory tract | 2 | 3 |
| 45 | 52 | F | AD | Olfactory bulb + tract (both) | 6 | 4 |
| 46 | 78 | F | AD | Olfactory bulb + tract | 2 | 0 |
|  |  |  |  | Olfactory bulb + tract | 3 | 0 |
| 47 | 79 | M | AD | Olfactory bulb + tract | 2 | 0 |
|  |  |  |  | Olfactory bulb + tract | 2 | 0 |
| 48 | 74 | M | OND | Olfactory bulb + tract | 4 | 0 |
| 49 | 79 | M | OND | Olfactory tract (both) | 2 | 0 |
| 50 | 81 | M | OND | Olfactory tract | 1 | 0 |

AON, anterior olfactory nucleus; LBD, Lewy Body disease; AD, Alzheimer's disease; OND, other neurodegenerative disease

## Notes

### Competing Interest Statement

The authors have declared no competing interest.

